# How chromatin-binding proteins direct distinct folding pathways of tetra-nucleosomes: Insights from coarse-grained simulations

**DOI:** 10.1101/2025.07.27.667063

**Authors:** Utkarsh Kapoor

**Author notes:** Corresponding Author: Utkarsh Kapoor – Department of Chemical and Biomedical Engineering, University of Wyoming, Laramie WY 82071, United States.

## Abstract

The dynamic coupling between chromatin organization and biomolecular condensates is governed by chromatin-binding proteins, yet the structural mechanisms by which these proteins modulate nucleosome interactions across spatial and organizational scales remain poorly understood. In this work, using high-resolution sequence-specific coarse-grained models combined with well-tempered metadynamics and parallel tempering, we investigate how heterochromatin protein 1α (HP1α) and a truncated construct of Polyhomeotic-like protein (tPHC3) influence the stability and folding pathways of tetra-nucleosomes, a minimal yet functionally informative chromatin model, under dilute and dense-phase conditions. While these proteins are known to drive distinct nuclear condensates their differential impact on chromatin topology and folding dynamics remains unclear. To address this, we ask: Do HP1α and tPHC3 stabilize or disrupt the canonical β-rhombus and α-tetrahedron nucleosome conformations? Are α-tetrahedron motifs transient intermediates or metastable states, and how do their prevalence and persistence depend on protein identity and phase context? To answer these questions, we analyze folding free energy landscapes, diffusion maps-based dimensionality reduced coordinates, and intermolecular interaction networks. Our simulations reveal that HP1α promote flexible, short-range nucleosome bridging and transient α-tetrahedron–like intermediates without stabilizing persistent structural basins. In contrast, tPHC3 stabilize α-tetrahedron–like motifs that scaffold folding toward the compact β-rhombus configuration characteristic of crystal-state tetra-nucleosomes. We find that this behavior arises from a context-dependent reorganization of multivalent SAM–linker interactions: in the absence of chromatin, self-association in dense phase conditions is mediated by linker–linker and linker– SAM contacts, while in the presence of nucleosomes, these linker-mediated interactions are suppressed, prompting compensatory SAM–SAM assembly. This reorganization highlights the essential role of SAM-mediated bridging in enabling long-range chromatin compaction. Together, our results demonstrate that under dense phase conditions α-tetrahedron–like motifs act as metastable intermediates rather than obligate folding end states, and their emergence depends critically on the identity of the chromatin-binding protein and their ability to mediate bridging. These insights offer a mechanistic framework for understanding how distinct architectural proteins encode topological preferences and remodel chromatin architecture across scales to support condensate formation and nuclear compartmentalization.

**Table of Contents Image:** 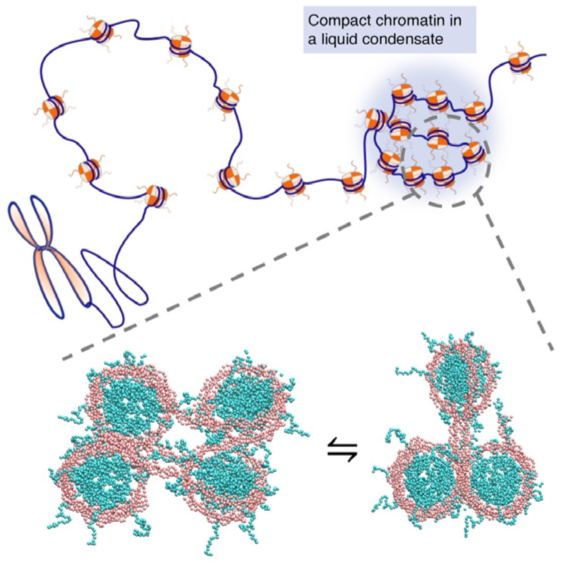

## 1. INTRODUCTION

The spatial organization of chromatin plays a critical role in gene regulation, epigenetic inheritance, and genome stability.(1–3) While nucleosomes serve as the fundamental packaging units of chromatin, higher-order chromatin folding remains highly dynamic and context-dependent, shaped by both intrinsic nucleosome interactions and extrinsic architectural proteins.(4–6) Over the past decade, there has been growing recognition that chromatin is not merely a static but a responsive scaffold whose structural states are coupled to the formation of biomolecular condensates – membraneless compartments that concentrate regulatory factors and modulate genome function through phase separation mechanisms.(7–13) Within this framework, short arrays of nucleosomes provide powerful model systems for dissecting chromatin folding principles. In particular, the tetra-nucleosome array has emerged as a minimal yet structurally informative unit that captures key features of chromatin compaction across spatial scales.(14–17) Crystallographic studies have revealed a canonical β-rhombus conformation stabilized by histone tail interactions and nucleosome stacking,(18,19) while cryo-EM and coarse-grained simulation studies suggest that alternative geometries, such as the α-tetrahedron, may also be accessible depending on the local environment and presence of binding partners.(14,16,20) Together, these results argue that intermediate states are not mere byproducts but can serve as functionally relevant, context-dependent architectural motifs. These findings further raise important questions about how chromatin-binding proteins, that form phase-separated condensates, modulate access to these different structural states.

In humans, Heterochromatin Protein 1 alpha (HP1α)(21–23) and Polyhomeotic-like protein (PHC3)(24,25) represent two archetypal chromatin-binding proteins with distinct structural architectures and biological roles. HP1α contains a chromodomain that recognizes H3K9me3 marks and a chromoshadow domain that mediates dimerization and bridging interactions, making it a hallmark component of constitutive heterochromatin and a model for dynamic chromatin compaction.(26–29) In contrast, PHC3 is a core member of Polycomb Repressive Complex 1 (PRC1) and features a sterile alpha motif (SAM) that drives oligomerization and long-range chromatin engagement through polymeric assembly – a mechanism critical for Polycomb body formation and the maintenance of facultative heterochromatin.(24,30–32) Notably, even truncated constructs of PHC3 that retain the SAM domain and flexible linker regions can undergo phase separation and compact chromatin *in vitro*,(25,33) making them tractable systems for controlled mechanistic dissection.

Despite increasing evidence linking these proteins to phase-separated condensates and chromatin reorganization, the molecular mechanisms by which these two proteins shape nucleosome– nucleosome interactions and influence folding pathways remain poorly resolved. HP1α, with its dimeric configuration and disordered hinge, serves as a model for flexible, transient bridging interactions that promote structural heterogeneity. In contrast, the SAM-mediated oligomerization of PHC3 imposes geometric constraints and directional assembly, potentially stabilizing specific chromatin topologies. These differences provide a compelling basis to interrogate how protein-specific dimeric state and interaction networks modulate chromatin folding and reorganize chromatin across spatial scales.

In this study, we ask the following key mechanistic questions: Are distinct structural motifs – such as the β-rhombus or α-tetrahedron – selectively stabilized by one protein over the other and whether these local folding intermediates are transient or metastable? How do their multivalent architectures and self-association interfaces influence short-range versus long-range chromatin compaction? And, to what extent do chromatin-binding proteins dynamically reorganize their interactions in response to the presence of nucleosomes, specifically nucleosomal DNA? We examine these questions through high-resolution sequence-specific coarse-grained molecular dynamics simulations enhanced by well-tempered metadynamics and parallel tempering. We focus on how pre-formed homodimers of HP1α and a truncated construct of PHC3 (tPHC3) differentially impact the folding of tetra-nucleosome arrays under both dilute and dense-phase conditions. Our analysis integrates structural, thermodynamic, and interaction network perspectives to illuminate the principles by which these architectural proteins modulate chromatin folding pathways and reorganize their interaction topologies in response to chromatin context. Our simulations reveal that HP1α homodimers promote flexible, short-range nucleosome bridging and transient α-tetrahedron–like intermediates, whereas tPHC3 homodimers stabilize α-tetrahedron motifs that scaffold progression toward the compact β-rhombus configuration. This behavior is accompanied by a reorganization of intermolecular interactions; in the presence of chromatin, tPHC3 shifts from linker-mediated self-association to SAM–SAM bridging, enhancing long-range chromatin compaction. Overall, our results suggest that α-tetrahedron–like structures serve as metastable intermediates whose stability and persistence are shaped by both protein identity and phase context. These findings inform our understanding of sequence-specific chromatin organization and shed light on how different classes of chromatin modifiers guide epigenetic regulation through distinct structural modalities.

The remainder of this article is organized as follows. We begin by examining the sequence determinants of HP1α and tPHC3 to identify the structural features that may underly their distinct modes of chromatin engagement. We then present a thermodynamic characterization of tetra-nucleosome folding across protein and phase contexts using Q–Rg free energy surfaces, revealing variations in chromatin compaction and structural stability. Next, we analyze folding trajectories using diffusion map–based dimensionality reduction to uncover the diversity of folding intermediates and the emergence of α-tetrahedron–like motifs as metastable states. We further investigate intermolecular contact networks to elucidate how multivalent protein–protein and protein–DNA interactions are reorganized in the presence of chromatin. We conclude with a discussion of the broader implications of our findings for chromatin architecture, condensate specificity, and nuclear organization. Finally, we describe the computational methodologies underlying our simulations and analyses.

## 2. RESULTS

To elucidate how chromatin-binding proteins modulate higher-order nucleosome architecture, we begin by examining the structural basis of the tetra-nucleosome and the sequence determinants of HP1α and a truncated construct of PHC3 (tPHC3) – two key regulators of nuclear condensates. While these proteins are implicated in chromatin organization and phase separation, the mechanistic principles that underlie their differential effects on chromatin topology remain poorly understood. In particular, we focus on how their distinct domain architectures, charge distributions, and self-association tendencies may influence chromatin folding, stability and flexibility at the molecular scale.

Figure 1a (top panel) shows the crystal structure of a tetra-nucleosome (PDB ID: 5OY7),(19) which serves as the core structural unit in our simulations. Prior structural studies have identified two major conformational states (also shown in Figure 1a): the more extended planar β-rhombus, where nucleosomes align in a diamond-like arrangement with fewer interactions,(16,17,20) and the compact α-tetrahedron, in which nucleosomes are arranged in a pyramid-like configuration stabilized by multiple inter-nucleosome contacts. These two configurations represent key intermediates in chromatin folding and provide a benchmark for probing how chromatin-binding proteins influence structural transitions. Our simulations aim to test whether pre-formed HP1α and tPHC3 homodimers stabilize, disrupt, or reorganize these hallmark nucleosome conformations. We assess how each protein modulates chromatin compaction, conformational flexibility, and structural transitions over time. By tracking distinct interaction patterns and folding trajectories induced by HP1α and tPHC3 homodimers, we aim to uncover molecular strategies by which multivalent protein interactions and bridging events alter chromatin organization.

**Figure 1.**
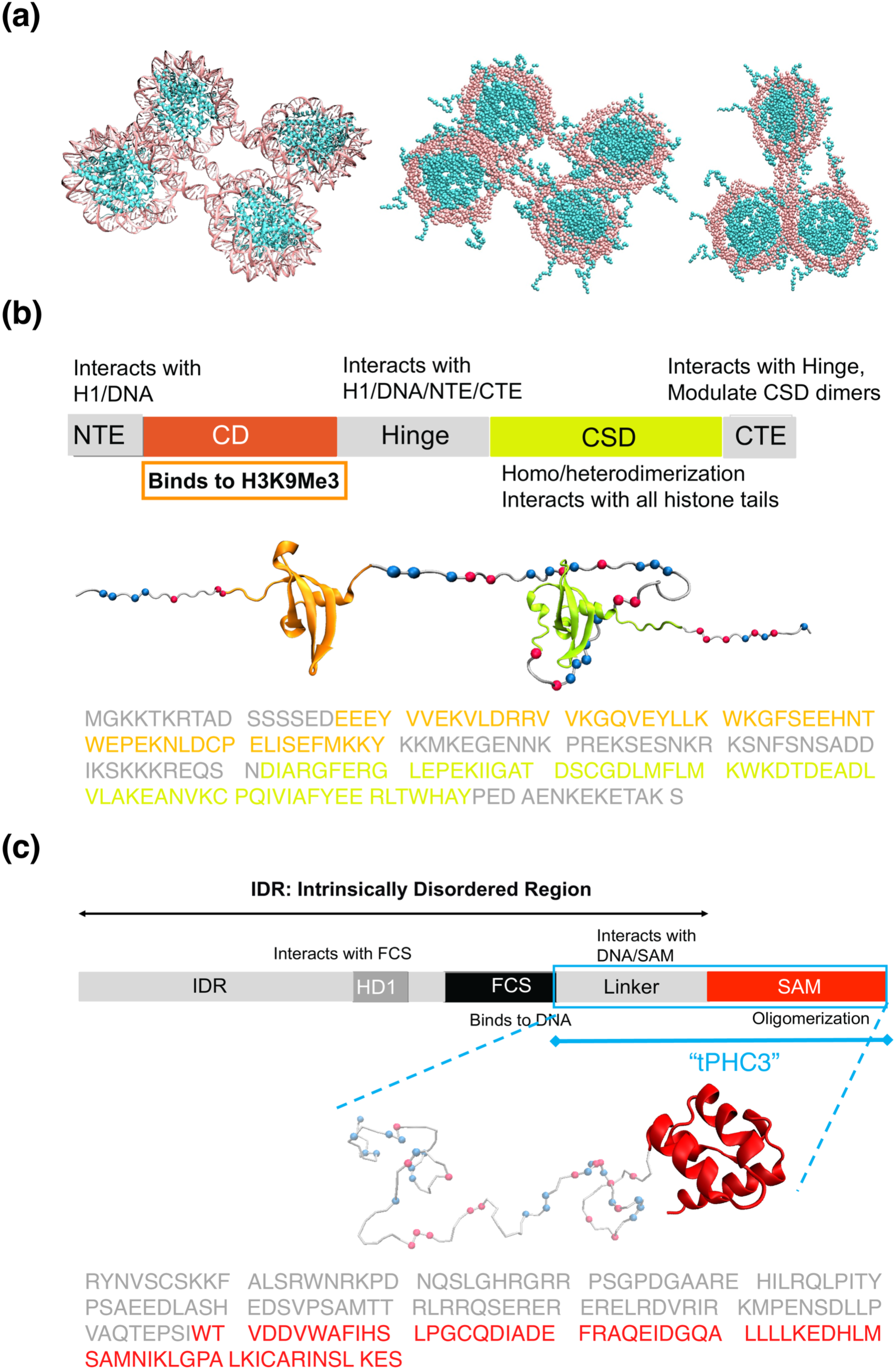
Structural and sequence features of tetra-nucleosome and chromatin architectural proteins. (a) The left panel shows the atomistic canonical crystal structure of the tetra-nucleosome (PDB: 5OY7), which serves as the model chromatin unit in our simulations. The middle panel illustrates the coarse-grained β-rhombus configuration, characterized by a planar arrangement of nucleosomes, while the right panel depicts the coarse-grained α-tetrahedron conformation, a compact pyramidal geometry formed by short-range nucleosome–nucleosome interactions. These two configurations represent hallmark folding states of short chromatin arrays and serve as reference conformations for assessing the impact of chromatin-binding proteins on folding pathways and structural transitions. (b) Domain architecture and sequence features of HP1α. HP1α is composed of an N-terminal extension (NTE), chromodomain (CD), a central hinge region, and the chromoshadow domain (CSD). These domains mediate key functions including H3K9me3 recognition, DNA binding, dimerization, and histone tail interaction. The full-length amino acid sequence is shown below, color-coded to match the domain schematic. Positively and negatively charged residues are marked with blue and red beads, respectively, to highlight the sequence-specific distribution of charges across domains. (c) Domain architecture and sequence features of PHC3, which includes an intrinsically disordered region (IDR), an FCS zinc finger, a flexible linker, and a sterile alpha motif (SAM) domain that mediates oligomerization. We use a truncated construct of PHC3 (tPHC3) in our simulations, retaining the key domains: SAM and the unstructured linker. The sequence of tPHC3 is displayed with domain-specific color coding, and charged residues are annotated with blue (positively charged) and red (negatively charged) beads to highlight electrostatic patterning across the protein. These domain-level annotations highlight the compositional and electrostatic contrasts between the disordered and structured regions of HP1α and tPHC3, setting the stage for interpreting their differential roles in chromatin organization.

### Structural basis for distinct chromatin engagement by HP1α and tPHC3

To interpret how HP1α and tPHC3 may influence nucleosome architecture, we first characterize the domain and sequence composition of HP1α and PHC3 (Figure 1b**–c**). HP1α comprises a disordered N-terminal extension (NTE), a chromodomain (CD) that specifically recognizes trimethylated lysine 9 on histone H3 (H3K9me3), a central hinge region that binds DNA, and a chromoshadow domain (CSD) that mediates homodimerization. In contrast, full-length PHC3 includes an intrinsically disordered region (IDR), an FCS zinc finger, a flexible linker, and a sterile alpha motif (SAM) domain that drives oligomerization through electrostatic pairing. While Figure 1c illustrates the full domain layout of PHC3, our simulations focus on a truncated construct tPHC3 consisting of the linker and SAM domain. This selection isolates the regions most directly involved in DNA binding and oligomer formation.

To gain a more quantitative perspective, we analyze the amino acid composition and charge distribution across full-length HP1α and tPHC3, as well as their key functional domains (Supporting **Figure S1**). HP1α is enriched in lysine (15.2%), glutamate (14.7%), and serine (7.85%), suggesting electrostatic versatility and a propensity for disordered regions. Its hinge domain, which mediates DNA binding, exhibits even greater polarization – dominated by lysine (26.8%), serine (17.1%), and asparagine (14.6%) – supporting a flexible, positively charged interface for interacting with chromatin. In contrast, the dimerizing CSD domain shows a more balanced composition (10.8% alanine, 10.8% glutamate, and 9.23% aspartate), indicative of a compact and globular fold. Charge-based descriptors reinforce these distinctions: the hinge domain has a net charge of +7 and a Sequence Charge Decoration (SCD)(34,35) parameter of +1.64, indicating strong local clustering of positive charges, while the full-length HP1α exhibits a net charge of –3 and an SCD parameter of –1.06, reflecting more dispersed charge distribution. These features align with previous reports of HP1α forming dynamic condensates and mediating nucleosome interactions through weak multivalent contacts, though the spatial extent and structural consequences of these interactions on chromatin remain incompletely understood.

In contrast, tPHC3 exhibits sequence features suggestive of different interaction modalities. The overall composition includes 11.0% arginine, 10.4% leucine, and 9.25% serine, blending hydrophobic and charged residues. The SAM domain, which mediates oligomerization, is rich in leucine (13.8%), alanine (10.8%), and aspartate (9.23%), supporting both hydrophobic core formation and electrostatic pairing at its multimeric interfaces. Although the SAM domain favors hydrophobic packing, its oligomerization interface involves multiple charged residues, underscoring the dual role of hydrophobic and electrostatic forces in assembly. The adjacent linker domain is particularly charged, with 15.7% arginine, 11.1% serine, and 9.26% glutamate. Charge metrics reflect this clustering: the linker alone carries a net charge of +5 and an SCD parameter of –0.74, while the linker–SAM construct shows a near-neutral net charge of +1 but an SCD parameter of –1.96. These profiles suggest the presence of oppositely charged patches that may contribute to DNA engagement or recruitment into biomolecular condensates.

Together, these compositional and biophysical distinctions suggest that HP1α and tPHC3 may engage chromatin through fundamentally different molecular strategies. While HP1α harbors a highly charged hinge and forms compact dimers, tPHC3 features spatially patterned charge and an oligomerizing SAM domain – indicating the potential for distinct modes of chromatin association. These molecular characteristics raise two central questions: how do HP1α and tPHC3 differ in the extent or persistence of chromatin bridging, such as short-versus long-range contacts; and how do these proteins reshape the conformational landscape of tetra-nucleosome arrays – stabilizing canonical α-tetrahedron or β-rhombus architectures, or favoring alternative metastable states? Our simulations directly probe these questions by examining how each protein perturbs nucleosome-nucleosome geometry, modulates structural flexibility, and reorganizes chromatin topology in ways that may underlie nuclear condensate function.

### Spontaneous Folding Landscape of the Tetra-nucleosome in the Absence of Binding Partners

To establish a baseline for chromatin folding in the absence of chromatin-associated protein interactions, we first examine the free energy surface (FES) of the protein-free tetra-nucleosome system from WTM-REMD simulations using the fraction of native contacts (Q) and the radius of gyration (R_g_) as structural order parameters (Figure 2). The crystallographic structure (PDB ID: 5OY7), corresponding to the compact β-rhombus configuration, maps to Q ≈ 1 and R_g_ ≈ 8.07 nm and serves as our reference for folded conformations. While our simulations recover this native basin, the FES also reveals a distribution of metastable intermediate states with Q values ranging from ∼0.6–0.9 and R_g_ values between 8.5–9.5 nm. These compact, yet partially unfolded conformations likely correspond to non-canonical packing topologies with disrupted nucleosome– nucleosome interfaces. At lower Q values (Q < 0.5), the FES becomes highly rugged, exhibiting broad sampling of extended conformations with R_g_ between 10–13 nm. These unfolded states represent a wide variety of loosely associated configurations lacking stable inter-nucleosomal contacts. We note that the negative correlation between Q and R_g_ restricts the data points to the lower bottom of the plane. The resulting folding landscape displays multiple accessible pathways connecting compact and expanded states, consistent with a dynamic ensemble of folded and misfolded chromatin geometries. The qualitative features of this FES landscape align with the study by Ding et al.,(17) which also utilized Q and R_g_ as collective variables and observed conformational plasticity in tetra-nucleosome arrays, implying convergence of the ensembles in our simulations in terms of the global dimensions.

**Figure 2.**
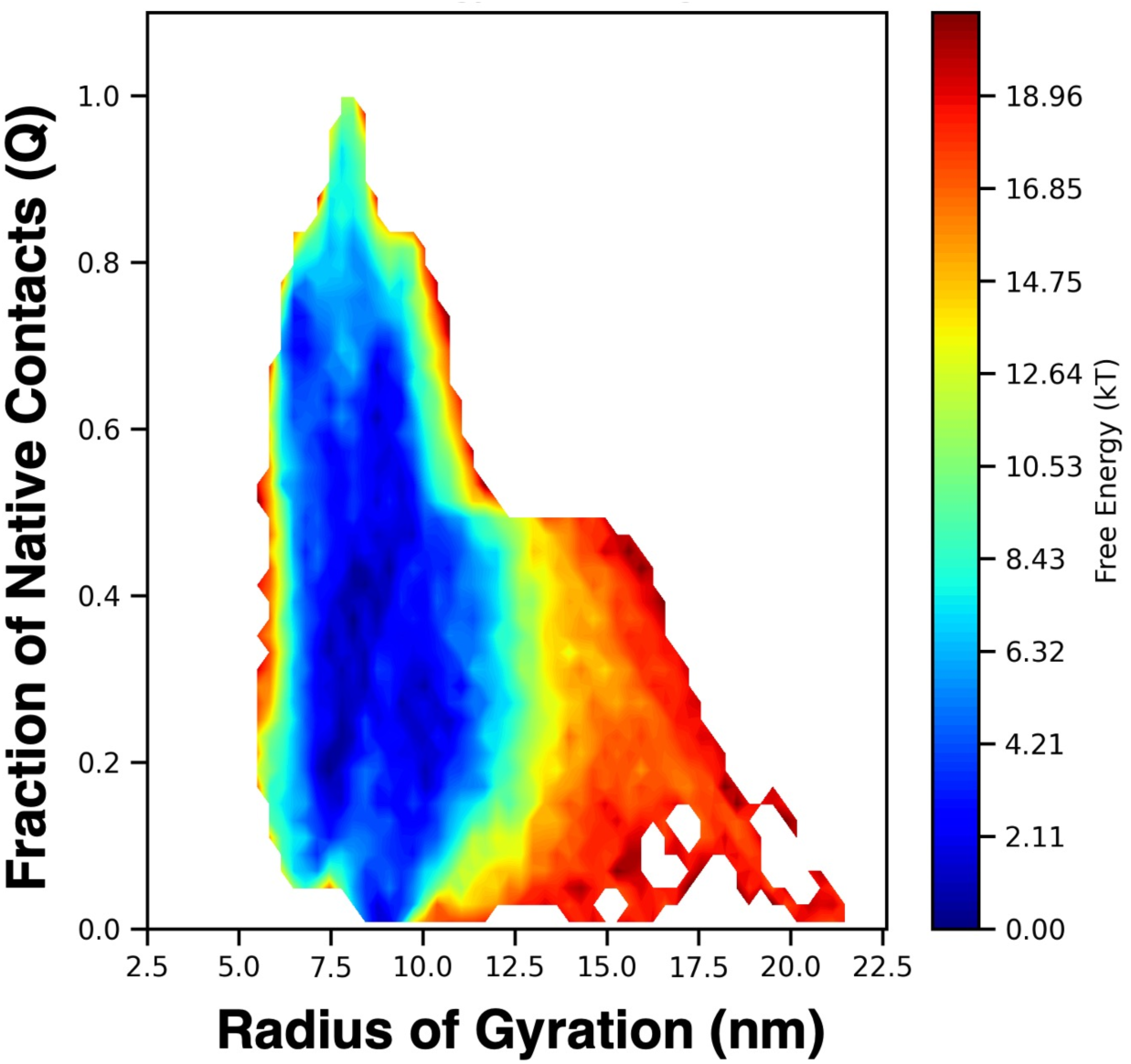
Free energy surface (FES) of the protein-free tetra-nucleosome system projected along the fraction of native contacts (Q) and radius of gyration (R_g_) structural order parameters. The FES reveals a well-defined native basin near Q ≈ 1 and R_g_ ≈ 8.07 nm, corresponding to the crystal structure (PDB ID: 5OY7), as well as several intermediate and unfolded states with reduced Q and expanded R_g_. The presence of metastable basins across a range of Q and R_g_ values highlights the structural plasticity of the tetra-nucleosome and provides a benchmark for assessing how chromatin-binding proteins modulate folding topology.

To better resolve conformational heterogeneity and folding pathways, we project demultiplexed trajectories from WTM-REMD simulations at 300 K of the binding partner free tetra-nucleosome onto a low-dimensional diffusion map constructed from pairwise structural contact similarities (Figure 3a). The resulting manifold, parameterized by diffusion coordinates ψ₁ and ψ₂, captures the slowest collective modes of structural variation across the ensemble. Coloring by root-mean-square deviation (RMSD) from the β-rhombus crystallographic structure (PDB ID: 5OY7) reveals a remarkably smooth and continuous gradient – ranging from green (low RMSD) to yellow (high RMSD) color – indicating a progressive structural ordering across the landscape. This organized color separation provides compelling visual support that ψ₁ correlates with global folding and compaction toward the β-rhombus–like conformation, while ψ₂ encodes lateral rearrangements among partially folded or misaligned nucleosome arrangements. The corresponding free energy surface (Figure 3c) reveals a rugged, funnel-like topology with a major low-free-energy basin near the origin and several peripheral basins extending along orthogonal directions. This central basin includes compact structures with low RMSD and Q ≈ 0.9–1.0, representing β-rhombus–like and near-native conformations. The extended arms of the funnel correspond to metastable intermediates with moderate to high RMSD, many of which feature distorted tetrahedral geometries or asymmetric packing. For instance, several sub-basins contain tri-nucleosome–like cores with the fourth nucleosome partially dissociated, rotated, or misaligned – states that also appear as intermediate basins in the Q–R_g_ projection (Figure 2). These alternative topologies highlight the presence of quasi-stable folding intermediates distinct from both canonical α-tetrahedron and β-rhombus geometries. Representative structural snapshots sampled across the ψ₁– ψ₂ plane (Figure 3c) illustrate this structural progression: from unfolded configurations at the funnel periphery, through compact yet asymmetric intermediates, toward well-packed, near-native states near the center. This suggests that folding proceeds not through a single discrete pathway but via multiple routes involving topologically distinct intermediates. Together, the diffusion map analysis reinforces the notion that the tetra-nucleosome ensemble includes a spectrum of conformations with varying degrees of order and compaction. In the absence of architectural proteins, the system explores both canonical folding states and non-native arrangements, raising key questions about whether HP1α or tPHC3 might stabilize specific topologies or redirect folding along distinct pathways.

**Figure 3.**
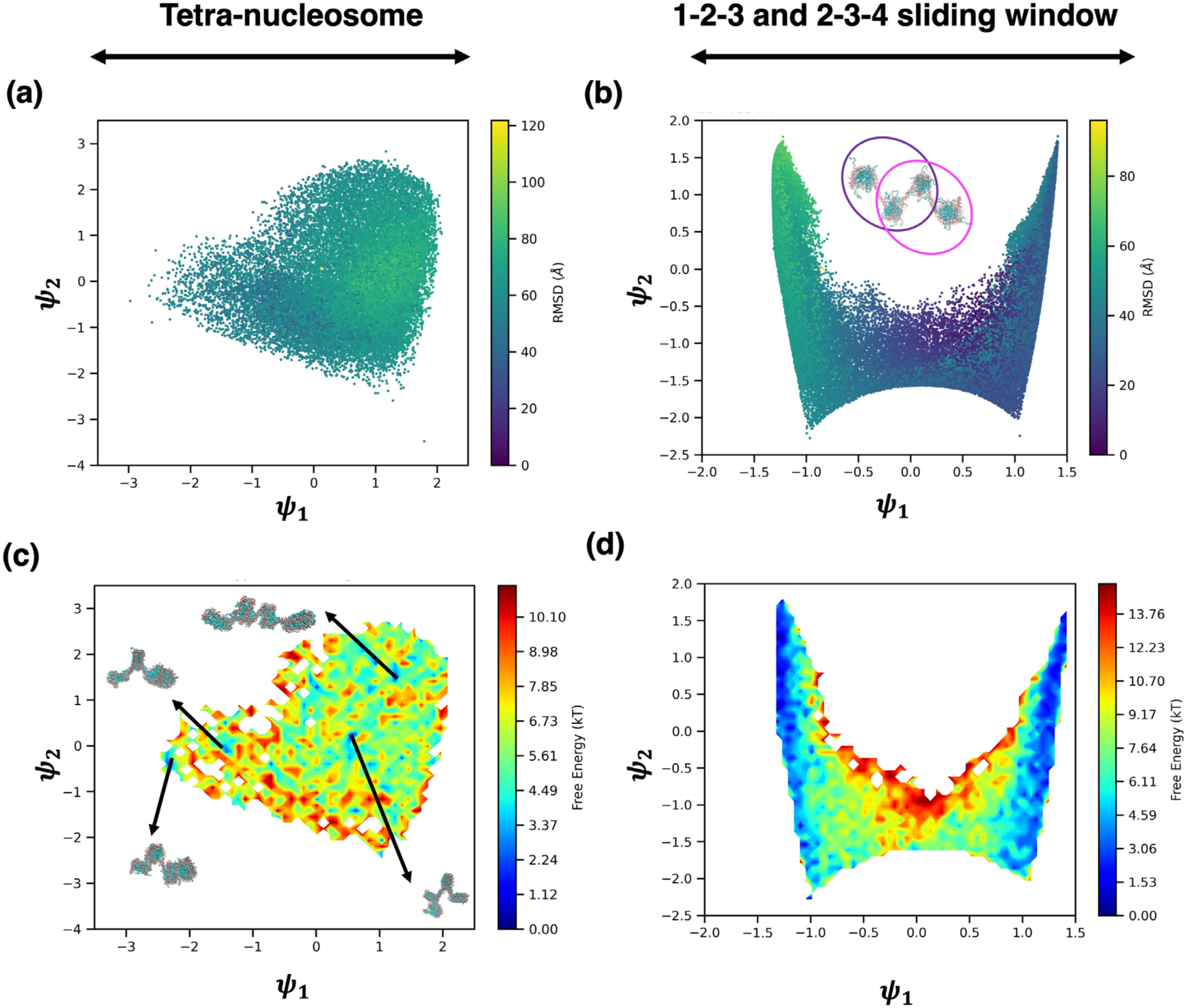
Diffusion map analysis of the protein-free tetra-nucleosome. (a) Diffusion map embedding of structural contact-based metrics for the full tetra-nucleosome system showing conformational spread along slow coordinates ψ₁ and ψ₂. (b) Sliding window analysis using 1–2– 3 and 2–3–4 nucleosome triads reveal a symmetric U-shaped diffusion landscape. (c) Free energy surface along ψ₁ and ψ₂ for the full tetra-nucleosome system, with representative structures illustrating transition paths between misfolded and folded basins. (d) Free energy surface of the sliding window data, showing the relative accessibility of different three-body configurations.

To further probe the internal topological evolution of the tetra-nucleosome, we apply a sliding window analysis to the demultiplexed trajectories from WTM-REMD simulations at 300 K by isolating sequential triads of nucleosomes i.e., Nuc1–Nuc2–Nuc3 or Nuc2–Nuc3–Nuc4, and analyzing their relative configurations using a diffusion map embedding (Figure 3b). The resulting manifold displays a distinct U-shaped topology, with two deep basins anchoring either end of ψ₁. These basins correspond to well-packed three-nucleosome arrangements that geometrically resemble a vertex of the α-tetrahedron. The central region of the map, in contrast, contains loosely packed and misaligned triads that sample disordered conformations. The corresponding free energy surface (Figure 3d) reveals that these α-tetrahedron–like substructures are thermodynamically accessible but separated by an ensemble of higher-energy intermediates. This suggests that during folding, the tetra-nucleosome may transiently visit locally ordered three-body motifs before globally assembling into a native conformation. Importantly, the duality of the U-shaped landscape – reflecting independent but symmetric basins for both triads – indicates that α-tetrahedron-like geometry can emerge from either half of the array, implying that partial folding may initiate cooperatively from either end.

To contextualize these transient local motifs, we also analyze the folding behavior of the tri-nucleosome system, which represents a simplified chromatin unit with three consecutive nucleosomes that adopts an α-tetrahedron geometry in its crystal structure. This system serves as a control for identifying local structural motifs but lacks the topological complexity introduced by a fourth nucleosome. The Q–R_g_ free energy surface (Supporting **Figure S2a**) reveals a compact and narrowly distributed basin centered near Q ≈ 0.9–1 and R_g_ ≈ 7.5–8.5 nm, closely matching the α-tetrahedron configuration of the crystal structure. The FES shows limited ruggedness and few accessible intermediate states, indicating that the tri-nucleosome predominantly adopts a well-packed, native-like geometry with little deviation. Unlike the tetra-nucleosome (Figure 2), which displays several partially folded basins at intermediate Q and elevated R_g_ values, the tri-nucleosome lacks such metastable extensions highlighting the importance of the fourth nucleosome in facilitating alternate packing arrangements. Diffusion map embedding of the tri-nucleosome trajectories (Supporting **Figure S2b**) reveals a distinctive triangular manifold in ψ₁– ψ₂ space. This triangular topology reflects constrained structural variability, with the apex of the triangle corresponding to compact, α-tetrahedron–like configurations and the sides representing minor geometric distortions, such as hinge-like rearrangements at the nucleosome–nucleosome interfaces. The absence of bifurcated branches or disconnected regions suggests a largely single pathway folding process with limited flexibility. The corresponding free energy landscape (Supporting **Figure S2c**) confirms dominant basins localized near the apex of the triangle, with no evidence of distinct alternate basins or unfolded intermediates. These features contrast sharply with the diffusion map obtained from tetra-nucleosome trajectories under the sliding window approach (Figure 3b,d), where two distinct branches emerge corresponding to variations in local three-nucleosome triads. The divergence of these branches and their associated structural heterogeneity are not reproduced in the tri-nucleosome system, reinforcing the conclusion that the fourth nucleosome introduces geometric frustration, longer-range couplings, or cooperative effects that diversify the conformational ensemble.

Together, these comparisons underscore a key distinction: while the tri-nucleosome adopts α-tetrahedron–like geometries, it lacks the structural plasticity and intermediate state diversity observed in the tetra-nucleosome system. Importantly, our findings indicate that α-tetrahedron– like motifs do arise transiently during tetra-nucleosome folding, but they are not resolved as a stable end state, nor do they appear to mediate a dominant folding path toward the β-rhombus configuration. Instead, they occupy peripheral regions of the conformational landscape, suggesting they serve as transient intermediates rather than obligatory checkpoints. Based on the findings so far, while we cannot conclusively determine whether the α-tetrahedron lies on or off the dominant folding pathway to the β-rhombus crystal structure, we can strongly suggest that the presence of the fourth nucleosome introduces structural routes and intermediates inaccessible to the tri-nucleosome alone.

### Phase-dependent tetra-nucleosome folding with HP1α and tPHC3 binding partners

To understand how architectural proteins modulate the tetra-nucleosome folding landscape, we next analyze simulations of tetra-nucleosomes with homodimers of HP1α and a truncated construct of PHC3 (tPHC3) in both dilute and dense phase conditions (Figures 4 **and 5**). These systems reveal distinct effects on folding stability, structural heterogeneity, and intermediate formation, depending on both the binding partner and phase context.

**Figure 4.**
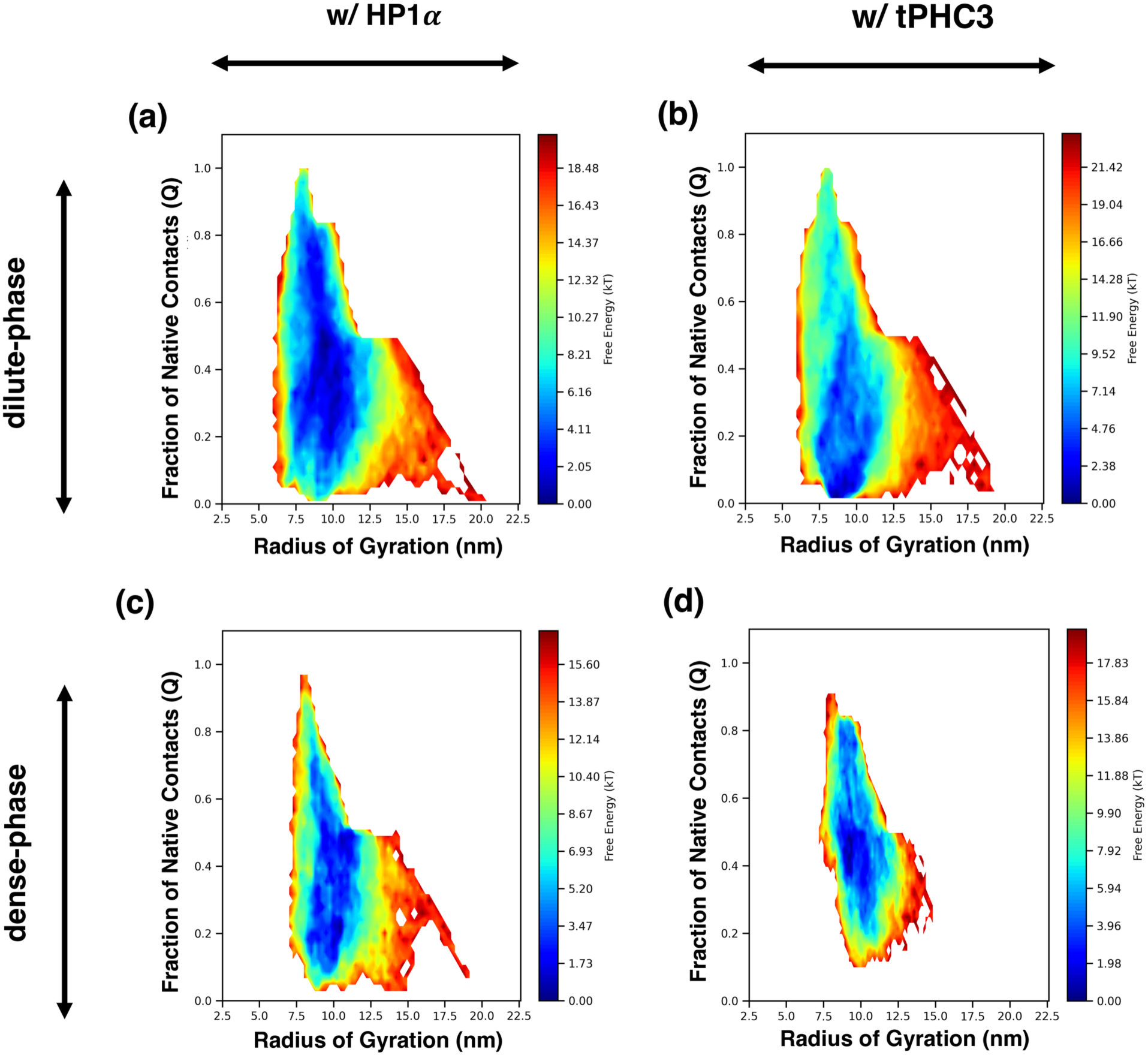
Free energy surfaces (FES) of tetra-nucleosome folding in the presence of HP1α and tPHC3 under dilute and dense phase conditions. FES are projected onto the Q and R_g_ structural coordinates, showing folding landscapes for (a) HP1α in dilute phase, (b) tPHC3 in dilute phase, (c) HP1α in dense phase, and (d) tPHC3 in dense phase. HP1α promotes folding under both conditions, with the folded basin shifting toward lower R_g_ and higher Q in the dense phase. tPHC3 exhibits a pronounced phase-dependent effect, showing minimal folding in the dilute phase and a sharp transition toward the native basin in the dense phase, consistent with enhanced compaction and stabilization of the folded state.

**Figure 5.**
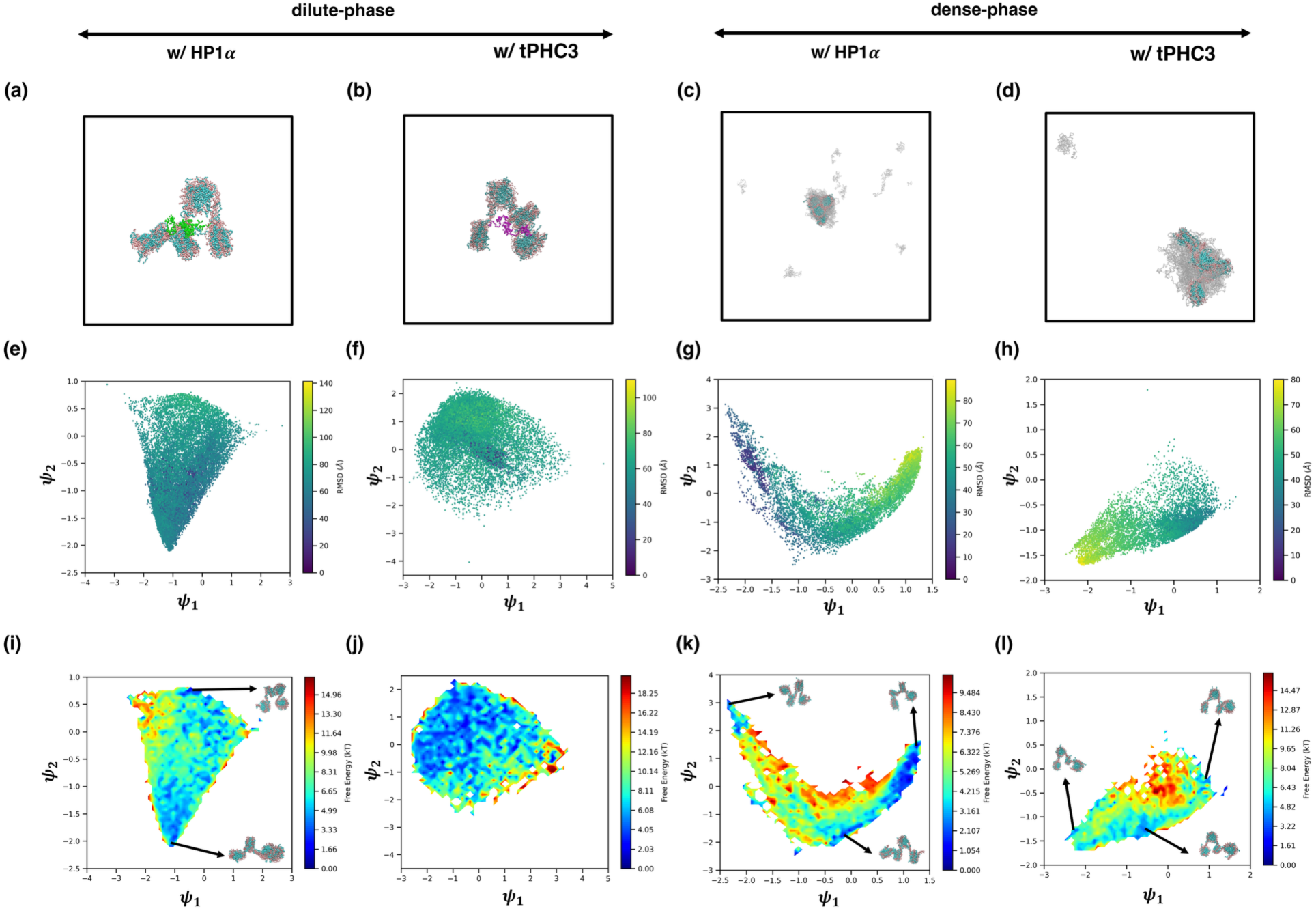
Diffusion map analysis and representative structures of tetra-nucleosome folding with homodimers of HP1α and tPHC3 in dilute and dense phases. (a–d) Representative structural snapshots from dilute and dense phase simulations. (e–h) Diffusion map projections (ψ₁, ψ₂) colored by RMSD to the native tetra-nucleosome structure. HP1α in both phases (e,g) shows organized landscapes with smooth RMSD gradients, indicating progressive folding. In contrast, tPHC3 (f,h) exhibits broader and more fragmented manifolds with multiple low-RMSD regions, suggesting heterogeneous folding pathways. (i–l) Corresponding free energy surfaces projected onto ψ₁–ψ₂ space. HP1α (i,k) favors continuous funnels leading toward native-like states, while tPHC3 (j,l) shows multiple quasi-stable basins. In the dense phase (l), several low-energy basins contain β-rhombus–like and α-tetrahedron–like structures (insets), highlighting the emergence of distinct intermediates enroute to compact chromatin architectures. The snapshots were rendered using VMD software.(36)

Free energy surfaces projected onto the Q–R_g_ space (Figure 4) show that HP1α homodimers promote compact conformations in both phases. In the dilute phase (Figure 4a), the FES exhibits a broad but shallow basin centered near Q ≈ 0.75–0.85 and R_g_ ≈ 10–12 nm. Upon transitioning to the dense phase (Figure 5c), this basin sharpens around Q ≈ 0.9 and R_g_ ≈ 8.5 nm, indicating a modest shift toward more cooperative folding. These restrained changes reflect HP1α’s binding mode – mediated largely through short-range bridging interactions between adjacent nucleosomes – which selectively reinforce local contacts without dramatically restructuring the global architecture. Even in the dense phase, the persistence of metastable arms in the FES implies that HP1α’s effect is additive and proximity-limited, enhancing local order rather than collapsing the ensemble into a uniquely folded state.

In contrast, tPHC3 homodimers induce a more dynamic and phase-sensitive reorganization of the folding landscape. In the dilute phase (Figure 4b), the FES shifts toward low-Q and expanded-R_g_ conformations, indicating limited folding and a predominance of disordered or partially assembled structures. However, the dense phase (Figure 4d) reveals a striking transition toward the compact native basin near Q ≈ 0.9 and R_g_ ≈ 8.5 nm, indicating a stronger compaction, enforcing a rigid topology, compared to the behavior observed with HP1α. This compaction trend is also reflected in monomeric tPHC3 simulations (Supporting **Figure S3**), which transition from a diffuse, low-Q distribution in the dilute phase (**Figure S3a**) to a sharply localized native-like basin in the dense phase (**Figure S3b**). Notably, the tPHC3 homodimer system achieves a broader but more connected FES basin in the dense phase (Figure 4d), suggesting enhanced accessibility to compact conformations. This enhanced folding is not merely a consequence of crowding but likely arises from tPHC3’s ability to form higher-order assemblies through its SAM domain. While monomeric tPHC3 displays some folding in the dense phase, pre-formed dimerization enables SAM–SAM interactions that seed oligomer formation across multiple nucleosomes. These oligomers may facilitate both short-range nucleosome stabilization and long-range bridging, thereby promoting cooperativity during folding. The resulting network-like architecture increases conformational connectivity and drives the system toward globally compact states. Thus, the structural influence of tPHC3 depends not only on phase environment but also on oligomeric state, which modulates its ability to organize nucleosomal contacts. Furthermore, unlike HP1α homodimer, which relies on short-range bridging interactions that stabilize compact states even in dilute conditions, tPHC3 homodimer appears to activate its bridging capacity primarily in dense environments. This likely reflects the requirement for multiple chains in proximity to nucleate SAM-mediated oligomerization. As a result, tPHC3 homodimer can bridge distal nucleosomes along the array, facilitating architectural reorganization that redefines the folding energy landscape. This distinction in interaction range – short for HP1α and long for tPHC3 – emerges as a key factor underlying their divergent compaction behaviors.

Diffusion map analysis further highlights differences in conformational heterogeneity and folding pathways across conditions. To this end, we project tetra-nucleosome ensembles onto low-dimensional diffusion maps constructed from pairwise contact similarities (Figure 5) and local tri-nucleosome sliding windows (Supporting **Figure S4**). Together, these analyses reveal how HP1α and tPHC3 homodimers modulate the global and local topologies of the folding landscape in a phase-dependent manner.

To aid visual interpretation, Figures 5a**–d** show representative snapshots from the dilute and dense phase simulations for HP1α and tPHC3 homodimers, respectively, illustrating the overall packing differences and system-scale organization across phase contexts. In the dilute phase, HP1α homodimers (Figure 5e) generates a moderately structured ψ₁–ψ₂ manifold with a dominant basin and several peripheral arms, consistent with a funnel-like organization toward native-like states. Structural snapshots from different basins and root-mean-square deviation (RMSD)-colored projections displaying a clear gradient from native-like (green) to disordered (yellow) conformations, suggests continuous folding progression through metastable intermediates, confirming the presence of compact, near-native geometries with minor topological variation, consistent with HP1α promoting cooperative folding without enforcing a rigid topology. These features are consistent with HP1α stabilizing compact configurations via short-range interactions. In contrast, tPHC3 homodimers in the dilute phase (Figure 5f) populates a broader and more dispersed ψ₁–ψ₂ space, lacking directional progression or a dominant folding basin. The corresponding free energy surface (Figure 5j) is diffuse and shallow, with elevated ruggedness and no clear energetic bias toward native conformations. RMSD coloring shows poorly resolved structural, ordering suggesting partial compaction with loosely associated nucleosomes. This pattern indicates structural frustration or incomplete folding. These findings also are consistent with diffusion map analysis of the monomeric tPHC3 system under dilute-phase conditions (Supporting **Figure S4a,e**), which shows an even more diffuse ψ₁–ψ₂ manifold and poorly resolved basins.

In the dense phase, both proteins promote compaction but via distinct topological and energetic features. HP1α homodimers (Figure 5g) yields a V-shaped diffusion map with a central folded basin and a curved arm of metastable intermediates, consistent with a moderately funneled free energy surface (Figure 5k). The RMSD-colored projection exhibits a structured gradient from disordered to native-like conformations, reinforcing the idea of a continuous folding route. Representative snapshots from the lower basins exhibit β-rhombus–like topologies with minor asymmetry, consistent with HP1α’s ability to reinforce compact states by bridging nearby nucleosomes through short-range interactions. In contrast, tPHC3 homodimers (Figure 5h) gives rise to a rugged and fragmented ψ₁–ψ₂ manifold with multiple quasi-stable basins, indicating the existence of several disconnected folding trajectories. The corresponding free energy surface (Figure 5l) shows prominent energetic barriers separating distinct minima, reflecting a heterogeneous ensemble of folding intermediates. Structural snapshots sampled from lower basins highlight β-rhombus–like arrangements and partially twisted configurations, consistent with tPHC3 promoting compaction through alternate topologies. The RMSD-colored diffusion map displays well-defined gradients and spatial organization, confirming enhanced folding relative to the dilute case. Furthermore, we note that ψ₁ behaves as a folding progress coordinate in both cases but with opposite directionality: in HP1α systems, native-like states occupy low ψ₁ values, while in tPHC3, folded conformations appear at high ψ₁ values. This inversion reflects differences in the dominant slow modes captured by diffusion map embedding and underscores the distinct folding topologies enforced by each protein. On the other hand, ψ₂ reveals critical differences in orthogonal heterogeneity. In the HP1α dense-phase system, ψ₂ spans a narrow range, consistent with a unified folding pathway through shallow intermediates. In contrast, the tPHC3 dense-phase system exhibits broad ψ₂ spread with discrete basins, indicating multiple structurally distinct folding trajectories and a more fragmented energy landscape. These features underscore the enhanced pathway degeneracy and topological diversity introduced by tPHC3-mediated compaction.

Moreover, this behavior observed in dimeric tPHC3 system is also echoed in the monomeric tPHC3 dense-phase system (**Supporting Figure S4c,g**), which, while still showing greater heterogeneity than HP1α, populates a more structured ψ₁–ψ₂ space than its dilute-phase counterpart. The ψ₁–ψ₂ manifold (Figure S4c) includes curved arms and localized density islands, and the free energy surface (Figure S4g) reveals shallow, spatially organized basins – many of which correspond to compact α-tetrahedron–like motifs. While these monomer-driven basins are less deep and connected than those in the dimer system, they still include α-tetrahedron–like intermediates, confirming that dense-phase conditions alone can promote local folding even without oligomerization. Altogether, the dense-phase folding behavior of tPHC3 reflects a hierarchical mechanism – where local structuring emerges from individual SAM-linker units, and higher-order compaction is amplified through multivalent bridging enabled by oligomerization. This stands in contrast to HP1α’s folding mechanism, which – despite also being dimeric – relies on short-range, stabilizing contacts that reinforce local interactions without driving topological transitions. These findings emphasize that it is not just multivalency, but the architecture of multivalent engagement, that governs how each protein reshapes the folding energy landscape.

To evaluate whether α-tetrahedron–like local packing acts as a dominant intermediate enroute to the β-rhombus configuration during tetra-nucleosome folding, we apply a sliding window diffusion map analysis to capture three-body structural motifs along the folding trajectory (Supporting **Figure S5**). This approach enables a focused assessment of tri-nucleosome geometries embedded within the larger four-nucleosome array, revealing whether local folding resembles known architectural motifs. In the dilute phase, HP1α homodimers produce a continuous ψ₁–ψ₂ distribution with mild enrichment in α-tetrahedron–like regions, but without a dominant basin indicative of persistent motif formation. This suggests that while HP1α may promote some level of local organization, these intermediates are transient and not the preferred structural scaffold in dilute conditions. By contrast, tPHC3 homodimers yields a fragmented landscape with sparse α-tetrahedron–like features, reflecting significant structural frustration and a lack of coordinated local folding. This behavior is mirrored in monomeric tPHC3 simulations (Supporting **Figure S4b,f**), where both the ψ₁–ψ₂ projection and free energy map reveal diffuse, weakly populated basins with no clear α-tetrahedron signature. In the dense phase, sliding window analysis reveals a notable shift. For HP1α homodimers, the ψ₁–ψ₂ landscape becomes more structured, and while β-rhombus–like local motifs become more prevalent, α-tetrahedron–like intermediates are still observed but do not dominate. The picture is different for tPHC3 homodimers, where the ψ₁–ψ₂ map fragments into multiple low free energy basins, several of which correspond to well-packed, α-tetrahedron–like geometries. This is consistent with the structural snapshots extracted from the lower basin of the full-system diffusion map (Figure 5l), which visually confirm the emergence of α-tetrahedron–like intermediates along paths to the compact β-rhombus state. Furthermore, monomeric tPHC3 in dense phase (Supporting **Figure S4d, h**) also displays enhanced enrichment of α-tetrahedron motifs, though to a lesser extent and with reduced landscape connectivity. These results suggest that dense-phase conditions enhance the formation and stabilization of α-tetrahedron–like local structures, and that dimerization of tPHC3 (via SAM domain) amplifies this effect, possibly by enabling cooperative recruitment of distal nucleosomes into coordinated folding pathways. Altogether, the sliding window analysis supports the view that particularly under dense-phase conditions α-tetrahedron–like tri-nucleosome geometries emerge as key intermediates during tPHC3-mediated folding. While not exclusive, their recurrent presence in low free energy regions and along folding pathways – especially in the rugged landscape of tPHC3 – indicates they likely play a transitional role enroute to β-rhombus packing. These intermediates appear to scaffold local compaction events that, when stitched together by bridging interactions, facilitate the assembly of the full tetra-nucleosome into the final folded architecture.

### Reactive folding trajectories reveal pathway heterogeneity and the role of tPHC3 oligomerization

To dissect the folding mechanisms underlying tetra-nucleosome compaction, we extract a reactive subtrajectory from WTM-REMD simulations at 300 K that begin in disordered (unfolded) states and end in folded conformations. We analyze three systems – (1) the protein-free tetra-nucleosome, (2) the tetra-nucleosome partitioned in the dense phase of HP1α homodimers, and (3) the tetra-nucleosome partitioned in the dense phase of tPHC3 homodimers – to assess how different chromatin-associated proteins and phase contexts influence folding routes.

We first project these subtrajectories onto the Q–R_g_ space to visualize folding as a function of native contact formation and global compaction (Figures 6a**–c**). All systems trace broadly uphill pathways, beginning in high-R_g_, low-Q regions and converging toward compact, high-Q folded basins. However, key distinctions emerge in trajectory geometry. The protein-free system (Figure 6a) follows a relatively narrow, directed path, indicating spontaneous folding through a preferred thermodynamic route. HP1α homodimers (Figure 6b) promote a smoother, more continuous progression, with folding intermediates aligned along a structured Q–R_g_ ridge – suggesting a guided funneling mechanism supported by multivalent interactions. In contrast, tPHC3 homodimers (Figure 6c) generate a more dispersed trajectory, marked by stalls, lateral excursions, and bifurcations in Q–R_g_ space. This heterogeneity suggests a more rugged folding landscape that accommodates competing routes or intermediate states.

**Figure 6.**
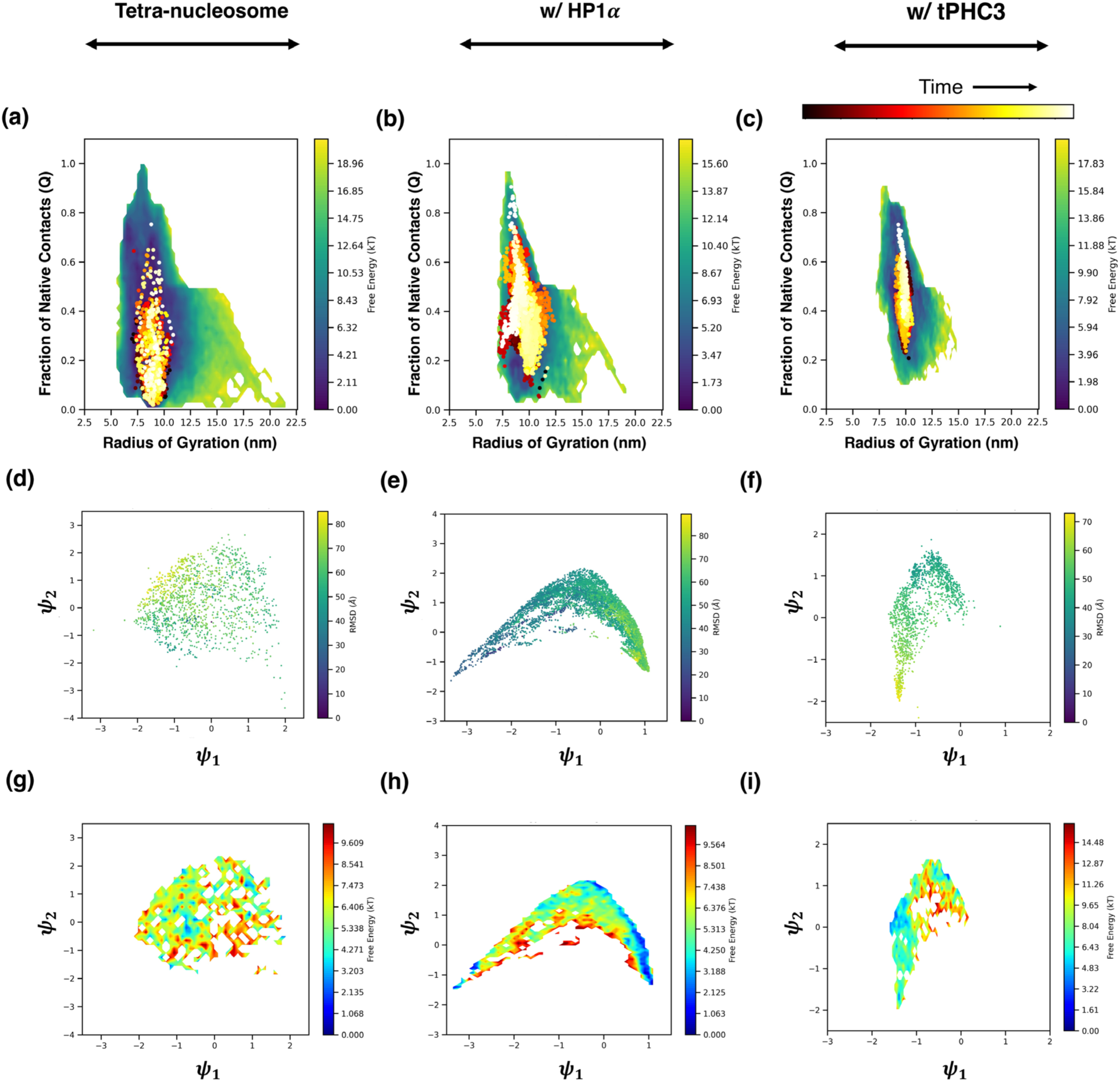
Reactive folding subtrajectories reveal condition-dependent heterogeneity and pathway diversity. (a–c) Q–R_g_ free energy surfaces for reactive subtrajectories that start from disordered unfolded and end in folded tetra-nucleosome conformations, shown for the protein-free system (a), HP1α-bound system (b), and tPHC3-bound system (c). Overlaid are time-colored points tracing folding progression (black → red → yellow). (d–f) Diffusion map projections (ψ₁–ψ₂) of the same subtrajectories, colored by RMSD from the native crystal structure (green ≈ low RMSD; yellow ≈ high RMSD). ψ₁ serves as an approximate folding coordinate, though its directionality varies between HP1α and tPHC3. (g–i) Corresponding free energy surfaces projected onto ψ₁–ψ₂ space, where blue indicates low and red indicates high free energy. Protein-free and HP1α systems exhibit relatively continuous folding pathways with single dominant basins, while tPHC3 displays a more fragmented landscape with multiple metastable intermediates. These differences in folding topology reflect the influence of chromatin-binding proteins on the conformational dynamics and energy landscape of nucleosome array folding.

To further resolve conformational diversity along these trajectories, we apply diffusion map embedding to the reactive segments using RMSD-based pairwise distances. Figures 6d**–f** show the resulting ψ₁–ψ₂ coordinates colored by RMSD from the native crystal structure, while Figures 6g**–i** present corresponding free energy surfaces. For the protein-free system (Figure 6d), the tetra-nucleosome trajectory progresses smoothly along ψ₁ with minimal variation along ψ₂, consistent with a relatively direct disordered-to-folded transition. The corresponding free energy surface (Figure 6g) displays a single, dominant basin and low ruggedness, reflecting a well-defined folding route. The HP1α-bound system (Figure 6e) also exhibits a largely monotonic decrease in RMSD along ψ₁, though the direction of structural progression is reversed compared to the tPHC3 system (ψ₁ increases rather than decreases with folding). Despite this inversion, the ψ₁ axis remains an effective progress variable. The HP1α free energy surface (Figure 6h) is more compressed along ψ₂ and exhibits fewer alternative minima, reflecting reduced conformational branching. This is consistent with HP1α’s role in promoting cooperative folding via short-range interactions that reinforce compact states without inducing topological reorganization. By contrast, the tPHC3-bound system (Figure 6f) traces a more fragmented and multidirectional path across ψ₁–ψ₂ space, with pronounced bifurcations and disjointed arms. These features are mirrored in the corresponding free energy surface (Figure 6i), which reveals multiple shallow basins separated by visible barriers. This pattern suggests the existence of structurally distinct, metastable intermediates and folding routes, indicative of a rugged energy landscape and reduced kinetic coherence. These findings extend the trends observed in the full ensemble folding landscapes (Figure 5), where tPHC3 promoted compact yet topologically diverse conformations, in contrast to the more funneled and cohesive manifolds of HP1α. It is noteworthy that overall ψ₁ serves as a folding progress coordinate. Its directional consistency across global and reactive folding manifolds suggests that ψ₁ robustly encodes the dominant slow mode of folding, albeit with protein-specific orientation due to differences in interaction topology and compaction routes. ψ₂, by contrast, encodes pathway branching and topological diversity, and its spread serves as a proxy for intermediate-state heterogeneity.

To further probe the molecular origins of this heterogeneity in dimeric tPHC3 system, we compare these results with the monomeric tPHC3 system under dense-phase conditions (Supporting **Figure S6**). The reactive subtrajectory for this system (Figure S6a) proceeds from high-R_g_, low-Q configurations to moderately compact structures. However, the ψ₁–ψ₂ embedding (Figure S6b) remains diffuse, with poorly connected arms and a disjointed progression. The corresponding free energy surface (Figure S6c) is shallow and fragmented, lacking a dominant funnel toward the native state. Notably, while the monomeric system does access native–like structures, the absence of deep, connected basins highlights the limited ability of individual tPHC3 units to drive hierarchical compaction. This contrasts sharply with the dimeric tPHC3 system (Figure 6f), where oligomerization enables multivalent bridging and access to multiple folding routes. Together, these comparisons emphasize the functional role of tPHC3 oligomerization in promoting higher-order organization and compensating for structural frustration at the local level.

Altogether, these reactive trajectory analyses reinforce the conclusion that tetra-nucleosome folding is kinetically heterogeneous and condition-dependent. Protein-free and HP1α-assisted folding proceeds via relatively coherent routes toward compact states, whereas tPHC3-bound folding navigates a broader ensemble of partially folded intermediates. Oligomerization is critical to tPHC3’s folding behavior: while individual SAM-linker chains support limited compaction, only the dimeric form induces multiple folding routes and enhances access to compact topologies. These results underscore how folding mechanisms are shaped not just by phase environment or native contacts, but by the molecular architecture of the proteins that scaffold and organize chromatin-like assemblies. Notably, the observed differences in pathway diversity and topological ruggedness reinforce the idea that HP1α and tPHC3 may promote compaction via distinct interaction ranges – with HP1α favoring more localized stabilization of native-like geometries, and tPHC3 enabling broader reorganization through extended contacts. This distinction, while emergent from folding behavior, may reflect fundamentally different modes of chromatin remodeling encoded by their multivalent architectures.

### Intermolecular interactions reveal distinct binding modes and reorganization of HP1α and tPHC3 binding partners in the presence of tetra-nucleosomes

To mechanistically contextualize our folding and structural observations, we examine protein–DNA and protein– protein intermolecular interactions across phase conditions and protein states. Figure 7 summarizes the contact probabilities between HP1α or tPHC3 homodimers and DNA in dilute and dense phases, averaged across reweighted full trajectories from WTM-REMD simulations. Each heatmap reports contact frequencies between protein and DNA residues, with DNA laid out linearly along the x-axis. Vertical stripes on the DNA axis represent contact hotspots, corresponding to superhelical locations on the nucleosomal DNA, while four distinct gaps denote linker regions connecting the four nucleosomes. These superhelical location sites serve as spatial landmarks for mapping interaction locations.

**Figure 7.**
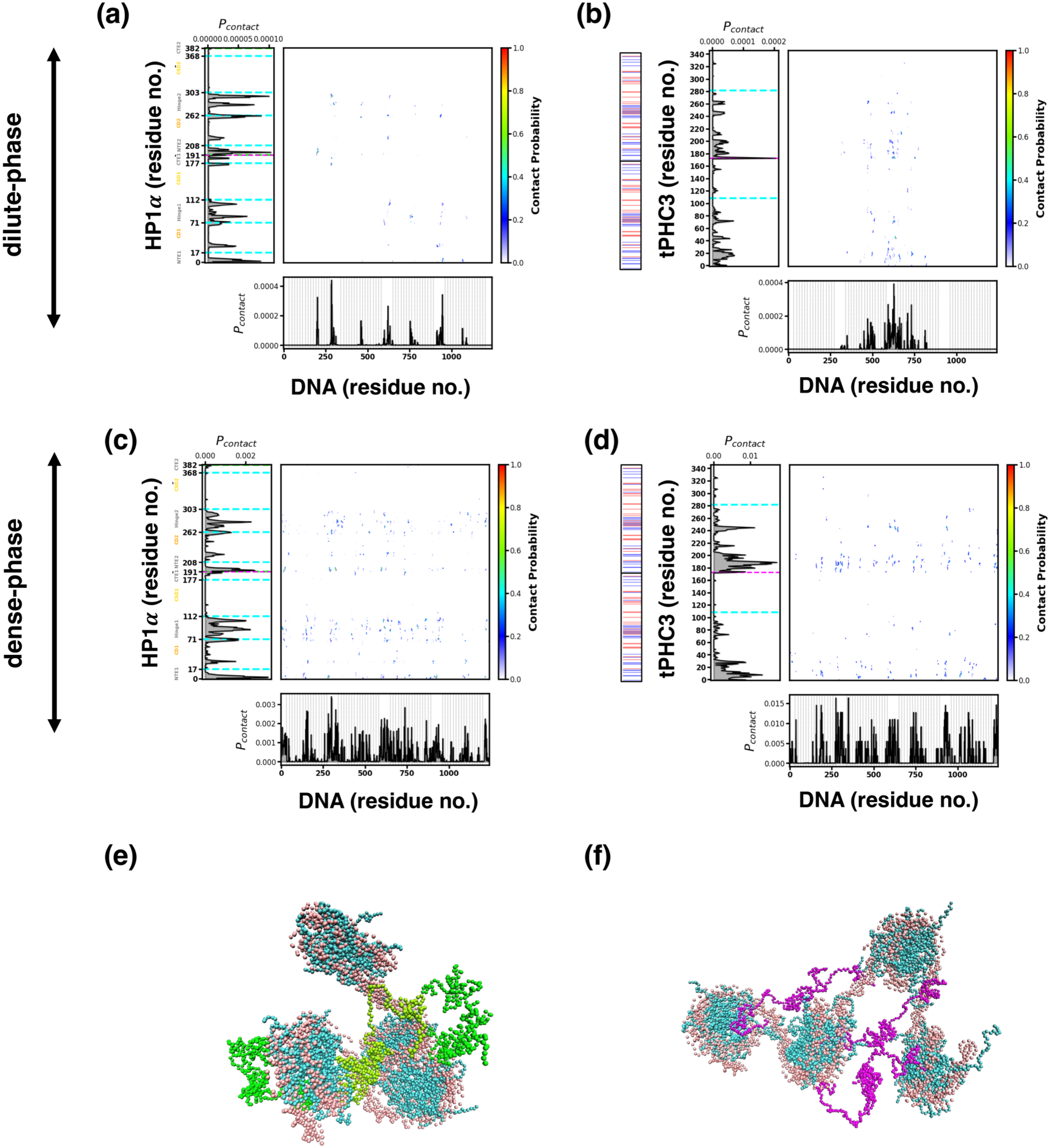
Protein–DNA interaction profiles and representative conformations of tetra-nucleosome systems with homodimers of HP1α and tPHC3 across phase conditions. (a–d) Contact probability maps between protein residues and DNA base pairs for HP1α (left panels) and tPHC3 (right panels) in the dilute phase (a,b) and dense phase (c,d), respectively, computed from reweighted WTM-REMD simulations. Vertical lines along the DNA axis indicate superhelical locations on the nucleosomal DNA, with four distinct gaps corresponding to linker regions between the four nucleosomes. Dashed lines along the protein axis delineate the boundaries between different domains of the protein. In the dilute phase, HP1α exhibits sparse but distributed interactions across nucleosomal and linker DNA, while tPHC3 displays restricted interactions localized to the central linker DNA near nucleosomes 2 and 3. Upon transitioning to the dense phase, both proteins increase their DNA engagement; HP1α maintains broad contacts with enrichment at dyad and linker regions, whereas tPHC3 shows enhanced, yet patchy interactions spanning multiple nucleosomes and superhelical locations, with minimal SAM domain involvement. (e, f) Representative simulation snapshots of the tetra-nucleosome in the dense phase in the presence of HP1α (e) and tPHC3 (f). DNA is shown in pink and histone core is shown in cyan, with HP1α shown in green and tPHC3 shown in magenta. For tPHC3 (f), linker-DNA engagement and SAM-domain bridging are evident, visually reinforcing interaction patterns observed in panel (d). The snapshots were rendered using VMD software.(36)

In the dilute phase, HP1α homodimers (Figure 7a) exhibit relatively sparse but spatially specific protein–DNA contacts, with pronounced enrichment in its hinge region – consistent with prior evidence identifying the hinge as a key DNA-binding motif. These interactions localize near the dyad and linker regions of nucleosomal DNA, spanning multiple superhelical locations and linker entry/exit points. This distribution suggests that HP1α homodimer is well-positioned to flexibly scan the chromatin fiber and transiently bridge adjacent nucleosomes, potentially stabilizing local folding intermediates without requiring extensive compaction. In contrast, tPHC3 homodimers (Figure 7b) show fewer overall contacts, which are narrowly concentrated around the central linker DNA between nucleosomes 2 and 3 of the tetra-nucleosome array. Notably, there is little to no SAM domain engagement with DNA under these conditions. The confinement of interactions to this central region, together with the absence of broader binding across superhelical locations, indicates limited capacity for scaffolded folding. However, this spatial targeting near the middle linker may reflect a poised configuration that enables long-range chromatin bridging and compaction, consistent with subsequent phase-dependent reorganization observed in dense-phase simulations.

Upon transitioning to the dense phase, both proteins exhibit enhanced DNA engagement, though with distinct interaction patterns. HP1α homodimers (Figure 7c) maintain broad DNA-binding behavior, with elevated contact frequencies near linker regions and nucleosomal dyads – consistent with its flexible, multi-point engagement and capacity to stabilize local chromatin contacts. In contrast, tPHC3 homodimers (Figure 7d) shows an overall increase in DNA binding, but with a more heterogeneous and spatially patterned distribution across superhelical locations. While all four nucleosomes participate in interactions, some superhelical location regions remain sparsely contacted, suggesting selective engagement rather than uniform DNA coating. Notably, contact density is enriched near the linker DNAs, which may serve as a focal point for higher-order architectural organization. Importantly, it is the linker domain of tPHC3 that interacts with DNA, and the SAM domain continues to show no detectable DNA interactions even in the dense phase, reinforcing the idea that tPHC3 compacts chromatin indirectly via SAM-mediated oligomerization rather than direct DNA binding. This asymmetric and discontinuous contact pattern reflects a distinct compaction strategy; wherein localized DNA contacts coupled with long-range protein– protein interactions drive structural reorganization of the tetra-nucleosome array. Figures 7e,f visually reinforces these mechanistic distinctions: HP1α homodimers decorate the chromatin surface through multiple, dispersed contacts, consistent with its short-range stabilization mechanism. In contrast, tPHC3 homodimers form extended linkages across distant chromatin segments, facilitated by SAM domain-mediated oligomerization. These visual distinctions complement our folding analyses, where HP1α promotes compact states via local bridging, while tPHC3 induces conformational diversity through extended topological reorganizations – highlighting fundamentally different modes of chromatin engagement.

In contrast, in the monomeric tPHC3 system, protein–DNA contact maps (Supporting **Figure S7a– b**) reveal a striking dependence on phase environment. In the dilute phase (Figure S7a), monomeric tPHC3 exhibits minimal DNA engagement, with contacts restricted to a single site and virtually no positional variability across the trajectory. This narrow interaction profile reflects an inability of the monomer to explore or stabilize diverse chromatin topologies. Upon transitioning to the dense phase (Figure S7b), monomeric tPHC3 shows markedly expanded DNA engagement, now contacting nearly all superhelical locations across the tetra-nucleosomal array. However, these contacts are broadly distributed and lack the domain-specific targeting observed in the dimeric system – for e.g., the SAM domain remains disengaged from DNA. This suggests that while crowding enhances non-specific contacts, it does not substitute for the architectural role of oligomerization in targeting and organizing chromatin. This behavior aligns with prior observations from folding analyses (Figures 5–6), which showed limited compaction and restricted pathway diversity in the monomeric system. Together, these findings reinforce the conclusion that oligomerization is critical for tPHC3’s functional chromatin engagement, enabling long-range bridging and structural modulation that remain absent in the monomeric state.

To further investigate how HP1α and tPHC3 self-associate and reorganize in response to chromatin context, we analyze intermolecular protein–protein contact maps in the dense phase, both in the presence and absence of tetra-nucleosomes. For the protein-only reference, we conduct unbiased phase coexistence simulations at 300 K and 100 mM salt, following protocols established in our prior work.(27,33,37)

In the presence of tetra-nucleosomes (Supporting Figure S8a), HP1α–HP1α contacts are broadly distributed and include both structured dimer interface interactions and transient associations among disordered regions. These interactions are consistent with its flexible binding architecture, which is known to support dynamic engagement with both DNA and protein partners. In prior studies of HP1α phase separation by us and others in the absence of nucleosomal arrays,(22,27) similar interaction patterns were observed – namely, enrichment of contacts at the hinge and chromodomain interfaces – highlighting HP1α’s ability to form multivalent, dynamic assemblies without adopting a rigid oligomeric interface. Our WTM-REMD results echo this behavior, with HP1α preserving flexible homotypic contacts while simultaneously engaging chromatin, suggesting a model in which short-range stabilization and transient bridging define its folding mechanism.

On the other hand, presence of tetra-nucleosomes leads to a marked reorganization of the protein– protein interaction landscape for tPHC3. In the protein-only dense phase (Supporting **Figure S9**), both monomeric and dimeric tPHC3 exhibit extensive linker–linker and SAM–linker interactions, forming a cohesive self-association network even in the absence of chromatin. However, upon introducing tetra-nucleosomes, these linker-mediated contacts are substantially diminished in both systems (Supporting **Figure S8b,c**), reflecting a functional reallocation of the linker region toward DNA engagement (as shown in Figure 8). This redistribution indicates that the linker cannot simultaneously support robust protein–protein and protein–DNA interactions under chromatin-condensed conditions. While this loss of linker–linker contacts is shared, the resulting oligomeric organization diverges between the dimeric and monomeric systems. In the dimeric case, the SAM– SAM interface is structurally pre-defined and constrained, preventing the emergence of additional cooperative interactions beyond those imposed by the modeled architecture. By contrast, the monomeric system (Supporting Figure S8c) undergoes significant reorganization: in the absence of pre-dimerization, SAM domains now form distinct and previously unobserved contacts with one another, compensating for the reduction in linker-mediated interactions. This reconfiguration reflects a context-specific shift in the oligomerization interface, emergent from chromatin-induced confinement. Notably, these emergent SAM–SAM contacts appear to be stabilized primarily through hydrophobic and aromatic interactions. These features enable SAM domains to form cohesive assemblies even in the absence of canonical head-to-tail alignment, mediated by charged residues, seen in crystallographic structures (Supporting **Figure S10**). These findings underscore the conformational adaptability of tPHC3 and its capacity to reorganize multivalent interfaces in response to environmental cues, including crowding and DNA engagement.

Altogether, these comparisons reveal that HP1α and tPHC3 employ fundamentally distinct modes of self-association. HP1α maintains a diffuse and resilient interaction pattern regardless of chromatin presence, reflecting its role in mediating short-range chromatin compaction. In contrast, tPHC3’s interaction network is highly sensitive to chromatin context: it reorganizes in response to DNA binding and relies on domain-specific cooperativity to scaffold multimeric structures. These behaviors align with our folding analyses, which showed that HP1α promotes coherent folding through flexible contacts, while tPHC3 enables broader topological variation via structured, multivalent scaffolding. Importantly, the capacity for interaction rewiring in tPHC3 underscores the functional importance of oligomerization: it allows the protein to tune its self-association mode to balance chromatin binding, structural organization, and phase behavior.

## 3. DISCUSSION

Our findings provide mechanistic insight into how chromatin-binding proteins, through their distinct multivalent architectures and interaction specificities, shape nucleosome–nucleosome interactions and guide chromatin folding into protein-dependent conformational states. Using enhanced sampling molecular dynamics and high-resolution sequence-specific coarse-grained models, we uncover how Heterochromatin protein 1 (HP1)α and a truncated construct of Polyhomeotic-like protein (tPHC3) differentially remodel the tetra-nucleosome folding landscape – modulating the balance between local packing motifs and long-range compaction pathways in ways that reflect their divergent roles in heterochromatin and Polycomb-associated domains.

A central result of this study is the context-dependent emergence of α-tetrahedron–like motifs (i.e., tri-nucleosome-like geometries) as metastable intermediates, rather than terminal states, along folding pathways. These motifs are favored to different extents depending on the protein and phase environment. Under dense phase conditions, while HP1α stabilizes transient α-tetrahedron–like intermediates and supports flexible, short-range chromatin bridging, tPHC3 – especially in its dimeric form – drives the system toward the compact β-rhombus configuration observed in tetra-nucleosome crystal structures. This occurs via a shift in interaction networks: linker-mediated contacts are repurposed upon chromatin binding, leading to the emergence of compensatory SAM– SAM interactions that promote longer-range compaction. These results underscore how chromatin architecture is not passively inherited from protein binding but is actively constructed through protein-mediated topological transitions. Importantly, these distinctions reflect not only differences in protein sequence and oligomeric state but also in their responsiveness to chromatin context. The reorganization of tPHC3’s SAM–linker network upon nucleosome engagement highlights a broader principle: chromatin-binding proteins can dynamically retune their interaction modes to accommodate environmental constraints, enabling selective stabilization of architectural motifs that drive phase-specific chromatin organization.

Our findings also raise broader implications for chromatin regulation across genomic scales. The ability to shift between transient intermediates and persistent packing geometries provides a potential mechanism for modulating chromatin accessibility, stiffness, and compaction in response to cellular cues. The differential preference for short-range versus long-range bridging interactions between HP1α and tPHC3 suggests that architectural proteins can encode not only condensate identity but also local chromatin topology and flexibility – parameters likely to be critical for controlling transcriptional states, epigenetic inheritance, and genome compartmentalization.

From a further broader perspective, our results raise new questions about how different chromatin-binding proteins partition spatially and functionally within the nuclear environment. Do combinations of proteins yield hybrid architectures with emergent folding pathways? Can sequence-dependent differences in DNA accessibility or nucleosome spacing shift the balance between α-tetrahedron and β-rhombus conformations? What roles do mechanical forces, histone modifications, or chromatin-associated RNAs play in tuning these transitions across nuclear compartments? How do local folding preferences propagate across larger chromatin domains to influence higher-order structure? These questions point toward a more nuanced view of chromatin as a responsive and reconfigurable scaffold – its organization shaped not only by local sequence and binding affinity but by the dynamic interplay of architectural proteins with multivalent, context-sensitive binding behavior. The identification of α-tetrahedron–like intermediates as common waypoints enroute to distinct folded states suggests a unifying motif in chromatin organization, one that may be selectively stabilized or bypassed depending on protein context.

Taken together, this work advances a mechanistic framework for understanding how nucleosome-level folding is tuned by the architectural features and interaction logic of chromatin-associated proteins. By linking multivalent binding behavior to folding trajectories and metastable motifs, our results illustrate how chromatin can serve as a responsive scaffold – its structure and dynamics shaped by the specific features of its protein partners. Moving forward, our framework can be extended to larger arrays, sequence-diverse regions, or mixed protein systems to probe how architectural complexity gives rise to spatially distinct chromatin domains. Ultimately, our findings lay the groundwork for deciphering how chromatin structure encodes regulatory logic – through folding intermediates, dynamic transitions, and multivalent scaffolding – at the intersection of molecular specificity and mesoscale nuclear architecture.

## 4. METHODS

### Model and simulation details for tetra-nucleosome folding

Motivated by the methodology of Ding et al.,(17) we use well-tempered metadynamics combined with replica exchange molecular dynamics simulations (WTM-REMD)(38–43) to sample the folding landscape of the of chromatin, using tetra-nucleosomes as a minimal yet functionally informative model. We perform all WTM-REMD simulations using LAMMPS package (Oct. 2020 version)(44) patched with the PLUMED (2.9.0 version)(45,46) enhanced sampling plugin for metadynamics sampling.

In the WTM framework, we use fraction of native contacts – a similarity-based order parameter – (𝑄𝑄) and radius of gyration (R_g_) as collective variables. Q quantifies the similarity between a given tetra-nucleosome configuration and the crystal structure, while R_g_ reflects global compaction. Thus, by biasing the simulations along Q and R_g_ – two structural descriptors that capture native-state formation – we directly examine tetra-nucleosome’s inherent folding landscape. Based on previous studies,(17) Q is defined as

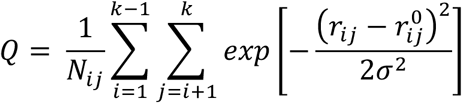

where 𝑖 and 𝑗 are indices of the nucleosomes, 𝑘 is the total number of nucleosomes in the array and is taken as 4 for tetra-nucleosomes. The sum runs over the total number of nucleosome pairs 𝑁*_ij_* and is taken as 6 for tetra-nucleosomes. 𝑟*_ij_* measures the distance between the center of nucleosome 𝑖 and 𝑗, and 𝑟*°_ij_* is the corresponding value in the crystal structure.(19) 𝜎 is taken as 2 nm.(17) R_g_ is computed as

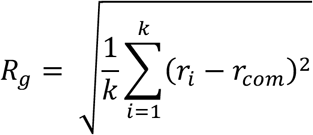

where 𝑟*_i_* is the geometric center of the 𝑖*^th^* nucleosome and 𝑟*_com_* is the center of mass coordinate for all nucleosomes in the array. We apply metadynamics bias along Q and R_g_ at 300 K using Gaussian kernels with widths of 0.01 and a height of 1.8 kJ/mol, with Gaussians deposited every 500 steps and a bias factor of 25. To avoid artifacts near the bounds of Q, we restrain it between 0.01 and 0.99 using harmonic walls with a spring constant of 10,000 kJ/mol and suppress bias beyond this range.(47,48) For the REMD component, we employ 16 temperature replicas, spaced geometrically from 300 K to 375 K, with replica swap attempts every 2500 steps. While WTM at 300 K enhances barrier crossing by introducing memory dependent potentials to penalize the system from revisiting the sampled region of phase space, REMD between 300-375K accelerates rare transitions via higher-temperature sampling.

To build a complete structure of the tetra-nucleosomes, we start with the coordinates from the crystal structure (PDB ID: 5OY7).(19) Since this structure lacks histone tail coordinates, we replace the histone proteins with those from the mono-nucleosome crystal structure (PDB ID: 1KX5)(49) to include histone tails, using UCSF Chimera.(50) We note that we keep the DNA sequence unchanged (see Figure 1). We combine the HPS-Urry protein model, a structure-based 𝐶_α_protein model,(51) and our recently developed 2-bead DNA model(37) to create a coarse-grained (CG) force field for describing the protein-DNA nucleosome complex, as we have done previously.(27,33,37) To enhance computational efficiency in CG simulations of the tetra-nucleosome, we model the core region of each nucleosome as a rigid body – grouping the ordered folded domains of the histone proteins with the 147 bp nucleosomal DNA. Positions and velocities of all CG beads within each rigid body are updated together, such that the rigid body translates and rotates as a single unit. Conversely, disordered histone tails and linker DNA remain flexible, with no restrictions on their conformational dynamics. Our partition of the rigid and flexible parts is informed by findings from prior studies.(17,52–56)

To simulate the nucleosome, we place the constructed CG tetra-nucleosome at the center of a large cubic simulation box with 120 nm dimensions. We relax the initial CG configuration using steepest descent energy minimization. Subsequently, we evolve the 16 geometrically spaced temperature replicas (300 K to 375 K) at 100 mM salt concentration for 2 μs each (total simulation time: 32 μs), using Langevin dynamics(57) in the NVT ensemble with a time step of 20 fs and a damping parameter of 10^5^ time steps. It is worth mentioning that though the CG DNA and protein models are parameterized using a 10 fs time step and 10^3^ damping parameter,(37,51) treating the nucleosome core as a rigid body removes high-frequency modes, permitting the use of a larger time step and damping parameter without compromising energy conservation or temperature control. We discard the first 0.1 μs of each replica as equilibration and save configurations every 5000 steps, yielding 19,000 snapshots per replica for analysis. Supporting **Figure S10a,e** shows the time evolution of Q and R_g_ at the 300 K replica. Evidently, the trajectory undergoes repeated folding and unfolding transitions through the entire RMSD and Q space, which indicates efficient conformational exchange between replicas and a uniform coverage of the phase space.

### Model and simulation details for modeling tetra-nucleosome with binding proteins

Motivated by our previous studies,(27,33,37) and to investigate how chromatin-associated proteins modulate the stability and folding of the tetra-nucleosome, we focus on two proteins implicated in chromatin organization through phase-separated condensates: Heterochromatin Protein 1 alpha (HP1α) and a truncated construct of Polyhomeotic-like protein 3 (termed tPHC3 in this study).

As shown in Figure 1, HP1α is a multidomain protein composed of two highly conserved folded domains – the chromodomain (CD) and the chromoshadow domain (CSD) – as well as three intrinsically disordered regions: the N-terminal extension (NTE), hinge region, and C-terminal extension (CTE). In comparison, tPHC3 consists of a single highly conserved folded Sterile Alpha Motif (SAM) domain and one disordered linker region. Previous studies have shown that CSD– CSD interactions in HP1α facilitate homodimerization, while SAM–SAM interactions in tPHC3 promote oligomerization. To examine the implications of these phase-separating proteins on tetra-nucleosome folding, we simulate systems with pre-formed HP1α and tPHC3 homodimers.

We construct the full-length HP1α homodimer using the protocol from our earlier work.(27) Specifically, we model the CD and CSD domains using crystal structures (PDB IDs: 3FDT(58) and 3I3C(59), respectively) and connect the disordered regions (NTE, hinge, and CTE) using MODELLER.(60,61) To generate the canonical HP1α homodimer, we arrange the CSD domains to interact through their known α-helix binding interface (3I3C). Similarly, we build the tPHC3 homodimer by positioning two SAM domains to interact via a head-to-tail interface modeled after PDB ID: 4PZO,(62) and connect the disordered linker using MODELLER.(60,61) These SAM– SAM contacts are shown in Supporting **Figure S11a**. The charge distribution on the SAM domains defines the head-to-tail oligomerization interface. Notably, the SAM-SAM binding interface predicted by AlphaFold3(63) also aligns closely with that observed in PDB ID: 4PZO, as illustrated in Supporting **Figure S11b**.

Like histone proteins, we construct a single-bead representation of each amino acid using the all-atom 𝐶𝐶_0_ positions and represent HP1α homodimer and tPHC3 homodimer by treating the CSD-CSD domains and SAM-SAM domains topologically together. We model the folded domains of HP1α homodimer and tPHC3 homodimer as rigid bodies.

To simulate protein–tetra-nucleosome systems, we consider two environments. In the first, which represents the dilute phase, we place single chain of HP1α or tPHC3 homodimer with a tetra-nucleosome chain in a 120 nm cubic simulation box. In the second, representing the dense phase, we place 50 chains of HP1α or tPHC3 homodimers alongside one chain of tetra-nucleosome in the same sized box. Unlike HP1α, which is known to exist as a stable homodimer *in vivo*, the oligomeric state of PHC3 in condensates remains less well-characterized. To further probe the role of SAM-mediated oligomerization on chromatin folding, we simulate a system containing 100 monomeric tPHC3 chains (chosen to match the total number of CG beads in the dimeric system) alongside one tetra-nucleosome chain. This comparison allows us to disentangle the specific contribution of tPHC3 self-association from general crowding effects and nonspecific interactions, thereby assessing whether pre-formed dimers are necessary for modulating chromatin structure.

We minimize the initial configurations using the steepest descent algorithm to eliminate steric clashes. We then perform phase coexistence simulations using the WTM-REMD methodology described previously, with slight modifications noted as following. In the dilute phase systems, to promote binding and unbinding between protein and nucleosome while ensuring protein– chromatin proximity, we apply a harmonic wall restraining the protein center-of-mass within 3 nm of the tetra-nucleosome center-of-mass using a spring constant of 200 kJ/mol. Whereas, in the dense phase simulations, we reduce the number of temperature replicas from 16 to 8 (geometrically spaced from 300 K to 350 K) to optimize computational cost. We simulate each replica for 2 μs at 100 mM salt concentration and save snapshots every 10,000 steps. We discard the first 0.1 μs of each replica as equilibration resulting in 9,500 frames per replica for analysis. We select the 300–350 K temperature range based on the known phase diagrams of HP1α and tPHC3.(27,33) Supporting **Figure S10** also includes time evolution of Q and R_g_ at the 300 K replica for the three systems. All systems exhibit broad fluctuations and various transitions between low-and high-Q states, indicative of reversible folding/unfolding events. Both Q and R_g_ show no systematic drift and fluctuate around well-defined means, suggesting that the systems have reached quasi-equilibrium. The persistent sampling of a wide range of Q and R_g_ values over multiple microseconds is consistent with the free energy landscapes presented in Figures 2, 4 and S3 and supports the thermodynamic relevance of the sampled basins. Thus, we consider that our simulations are well-equilibrated and have converged in the thermodynamically significant portions of the folding landscape.

### Analyses

We quantify the free energy surface (FES) of the tetra-nucleosome using two collective variables, Q and R_g_, both of which are biased during the WTM-REMD simulations. To recover the unbiased two-dimensional probability distribution P(Q, R_g_), we employ the reweighting scheme introduced by Tiwary and Parrinello,(64) which accounts for all deposited biases along the Q and R_g_ coordinates. The free energy surface is then calculated as:

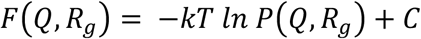

where kT is the thermal energy (the product of the Boltzmann constant and temperature), and C is an arbitrary additive constant.

To gain insights into the folding pathways, we analyze reactive trajectories by computing Q and R_g_ along continuous trajectories – i.e., trajectories that exchange temperature but remain continuous in time. We generate these time-continuous trajectories at 300 K by demultiplexing the parallel-tempered simulations. Within these demultiplexed trajectories, we explore the dynamical organization of the tetra-nucleosome folding process by applying diffusion map analysis(65) to the ensemble of reactive subtrajectories to uncover low-dimensional collective variables that capture the dominant conformational transitions during folding. This approach allows us to uncover low-dimensional pathways through which folding progresses or stalls, and to distinguish whether different proteins bias chromatin toward compact, misfolded, or intermediate metastable states. However, we note that in this study our focus lies primarily on the structural and thermodynamic features of different folding states. Therefore, we employ diffusion maps, which provide smooth, continuous, and structurally interpretable coordinates, rather than Markov state models(66,67) that are better suited for directly quantifying kinetic properties such as transition rates and fluxes. We perform this diffusion map analysis by adopting the DMAPS library(68) with in-house python scripts.

Following the methodology of Alvarado et al.,(16) we construct a diffusion kernel using a Gaussian weighting function applied to pairwise structural distances, where distance is computed using RMSD between translationally and rotationally aligned nucleosome coordinates in frames *i* and *j* of the trajectory, relative to the prototypical β-rhombus motif. The resulting diffusion map embedding reveals dominant modes of conformational rearrangement and enables us to move beyond traditional order parameters by clustering kinetically similar microstates in close proximity. To further provide physical interpretability of the diffusion map embeddings, we compute the RMSD of each contiguous stretch of 3 nucleosomes by passing a 3-nucleosome sliding window along the tetra-nucleosome array and comparing it to the prototypical α-tetrahedron motif. This comparative analysis reveals whether folding proceeds through canonical intermediates (e.g., α-tetrahedron) or diverges toward alternative pathways (e.g., bypassing β-rhombus), shedding light on how each protein modulates folding fidelity. We select the kernel bandwidth 𝜖𝜖 as 3.0 and diffusion map exponent α as 0.3 based on the median of the distance distribution to balance locality and global structure.(69) We compute the top three eigenvectors of the transition probability matrix and normalize the first two nontrivial diffusion components by the leading eigenvector, which corresponds to the stationary distribution. The resulting scaled coordinates are used to construct the diffusion map projections, which together account for the majority of short-time kinetic variance in the dataset. These coordinates serve as effective reaction coordinates that reveal dominant folding pathways, intermediate metastable states, and conformational bottlenecks. Overall, this dimensionality-reduction framework allows us to move beyond conventional order parameters and visualize the folding landscape in a kinetically meaningful way, where structurally and dynamically similar states cluster together. Motivated by prior studies,(16,70) we also construct the free energy surface over this intrinsic manifold to delineate both the metastable macrostates of the tetra-nucleosome and the interconversion pathways among them.

To further examine the implications of phase-separated condensates on tetra-nucleosome folding, within these demultiplexed trajectories, we also identify tetra-nucleosome subtrajectories that begin in the unfolded configuration and end in the correctly folded conformation, while partitioned within the condensate, and analyze intermolecular contacts among associated proteins HP1α or tPHC3, and between HP1α or tPHC3 and DNA. For contact analysis, we define two residues *i* and *j* as being in contact if they lie within 0.6 nm of each other.(27,37) By quantifying protein–DNA and protein–protein contacts, we identify how each protein facilitates or impedes nucleosome bridging, and how this correlates with folding success or trap formation. Unlike standard ensemble averaging, our use of reactive subtrajectories within phase-separated condensates enables direct correlation between folding outcomes and specific protein-mediated interactions in crowded nuclear-like environments. Furthermore, the time-resolved projection onto α-tetrahedron and β-rhombus RMSD landscapes allows us to identify whether each protein favors canonical folding routes or stabilizes off-pathway intermediates.

## AUTHOR CONTRIBUTIONS

Utkarsh Kapoor: Conceptualization; investigation; funding acquisition; writing – original draft, review and editing; formal analysis; visualization.

## DATA AVAILABILITY

Raw data and code are available upon reasonable request.

## SUPPORTING INFORMATION

Supplementary figures provide additional structural and energetic insights. These include residue-type compositions of HP1α and tPHC3 domains (Figure S1), tri-nucleosome folding landscapes (Figure S2), tetra-nucleosome folding in monomeric tPHC3 dilute and dense phases, sliding-window diffusion map analysis (Figure Figures S4–S5), reactive folding subtrajectories in monomeric tPHC3 dense phase (Figure S6), protein–DNA contact maps in monomeric tPHC3 dense phase (Figure S7), protein–protein interaction maps from WTM-REMD and unbiased simulations (Figures S8–S9), time evolution of metadynamics CVs: Q and R_g_ (Figure S10), and structural reference for SAM–SAM contacts from crystal and AlphaFold3 models (Figure S11). Together, these data complement and reinforce the mechanistic findings presented in the main text.

## ACKNOWLEDGMENTS

We acknowledge the financial support provided by the University of Wyoming, particularly through the NIH WY INBRE Pilot Award, which is supported by an Institutional Development Award (IDeA) from the NIGMS of the NIH under Grant #2P20GM103432. This research was partially completed with advanced computing resources generously provided by the University of Wyoming Advanced Research Computing Center (UW-ARCC), specifically the MedicineBow supercomputing cluster. Furthermore, we recognize the computing time on the Derecho system (doi:10.5065/qx9a-pg09) supported by the NSF National Center for Atmospheric Research (NCAR) at the NSF NCAR-Wyoming Supercomputing Center, sponsored by the National Science Foundation and the State of Wyoming.

