## Supplementary material for "How chromatin-binding proteins direct distinct folding pathways of tetra-nucleosomes: Insights from coarse-grained simulations": SI

***Utkarsh Kapoor***[1*]

*1. Department of Chemical and Biomedical Engineering, University of Wyoming, Laramie, WY 82071, United States*

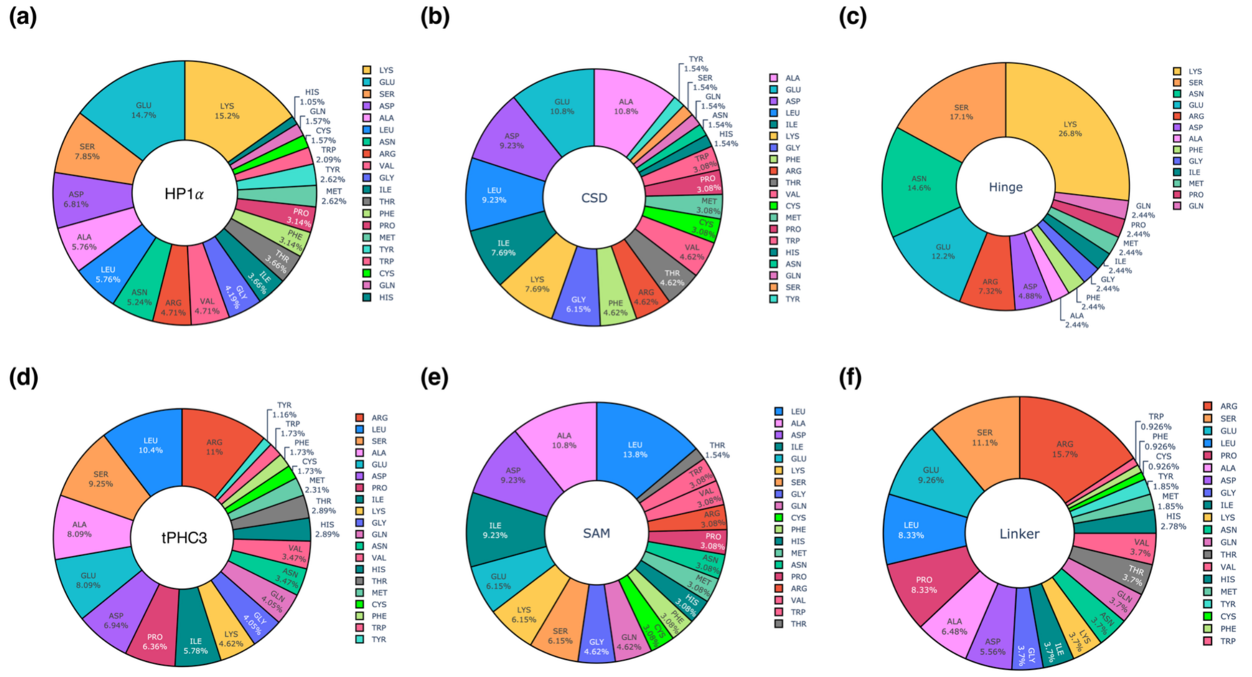


***Figure S1.*** *Amino acid composition profiles of HP1α and tPHC3 domains. Pie plots show residue-wise composition (% abundance) for: (a) full-length HP1α, (b) HP1α CSD domain, (c) HP1α hinge region, (d) tPHC3, (e) tPHC3 SAM domain, and (f) tPHC3 linker region. Compositional trends highlight functional specialization – HP1α hinge is enriched in lysine and polar residues, while CSD is more neutral. tPHC3 linker is rich in arginine and glutamate, while SAM domain favors hydrophobic amino acids. Together, these trends reflect sequence-driven mechanisms for DNA interaction, dimerization, or oligomerization.*

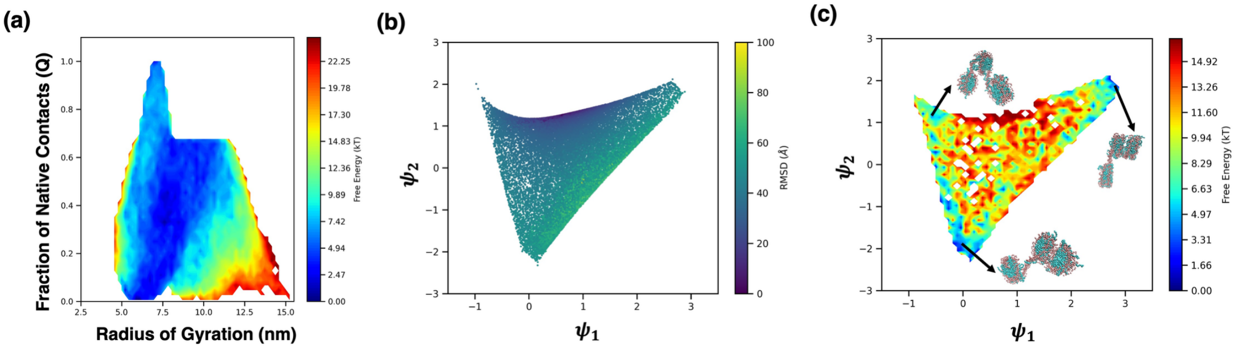


***Figure S2.*** *Folding landscape of the protein-free tri-nucleosome system. (a) Free energy surface along $Q$ and $R_g$ reveals a narrower conformational ensemble compared to the tetra-nucleosome. (b) Diffusion map embedding of tri-nucleosome structures.(c) Corresponding free energy surface highlighting the α-tetrahedron-like native conformation and limited sampling of unfolded states.*

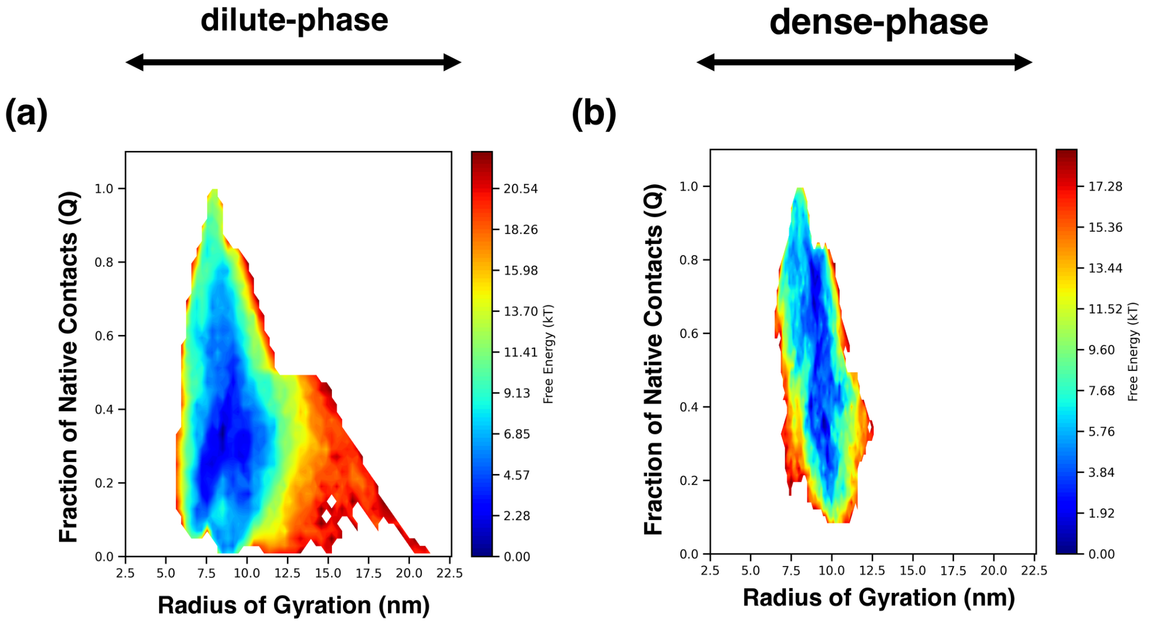


***Figure S3.*** *Free energy surfaces of monomeric tPHC3 in dilute and dense phases. The folding landscapes are projected onto the fraction of native contacts (Q) and radius of gyration (Rg), shown for (a) dilute phase and (b) dense phase. In the dilute phase, the FES is broad and shifted toward low-Q and high-Rg values, indicating predominantly disordered or partially folded conformations. In contrast, the dense phase exhibits a sharper and more focused basin centered, suggesting enhanced stabilization of compact, folded states. These results imply that phase context alone—independent of oligomerization—can modulate the conformational ensemble of tetranucleosome and facilitate folding.*

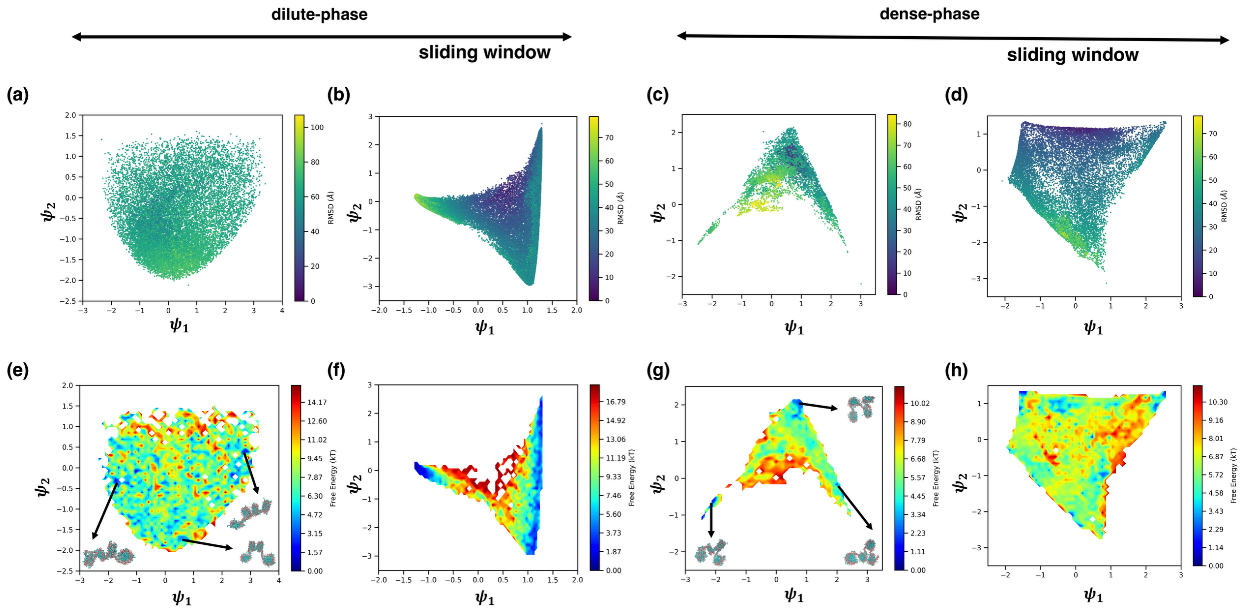


***Figure S4.*** *Diffusion map analysis of monomeric tPHC3 in dilute and dense phases, comparing full tetra-nucleosome and tri-nucleosome (sliding window) representations. (a–d) Diffusion map projections ($\psi_1$–$\psi_2$) for monomeric tPHC3 systems. Panels (a,c) depict the tetra-nucleosome–based projections in dilute (a) and dense (c) phases, while panels (b,d) show the corresponding tri-nucleosome (sliding window) projections for dilute (b) and dense (d) phases. The dense phase systems exhibit more confined and ordered manifolds, particularly in the sliding window view (d), suggesting emergence of local structural order not captured in the global tetra-nucleosome manifold. (e–h) Corresponding free energy surfaces projected onto $\psi_1$–$\psi_2$ coordinates. In the dilute phase, both tetra-nucleosome (e) and sliding window (f) maps show dispersed and fragmented basins, consistent with poorly ordered ensembles. In contrast, the dense-phase maps (g,h) display distinct low-energy basins. Notably, the sliding window dense-phase projection (h) shows compact basins enriched in α-tetrahedron–like geometries, indicating that local folding intermediates resembling α-tetrahedral packing may be stabilized even without dimerization. These results support the hypothesis that tPHC3 monomers can promote early-stage local structuring in dense environments, potentially seeding architectural transitions toward compact states.*

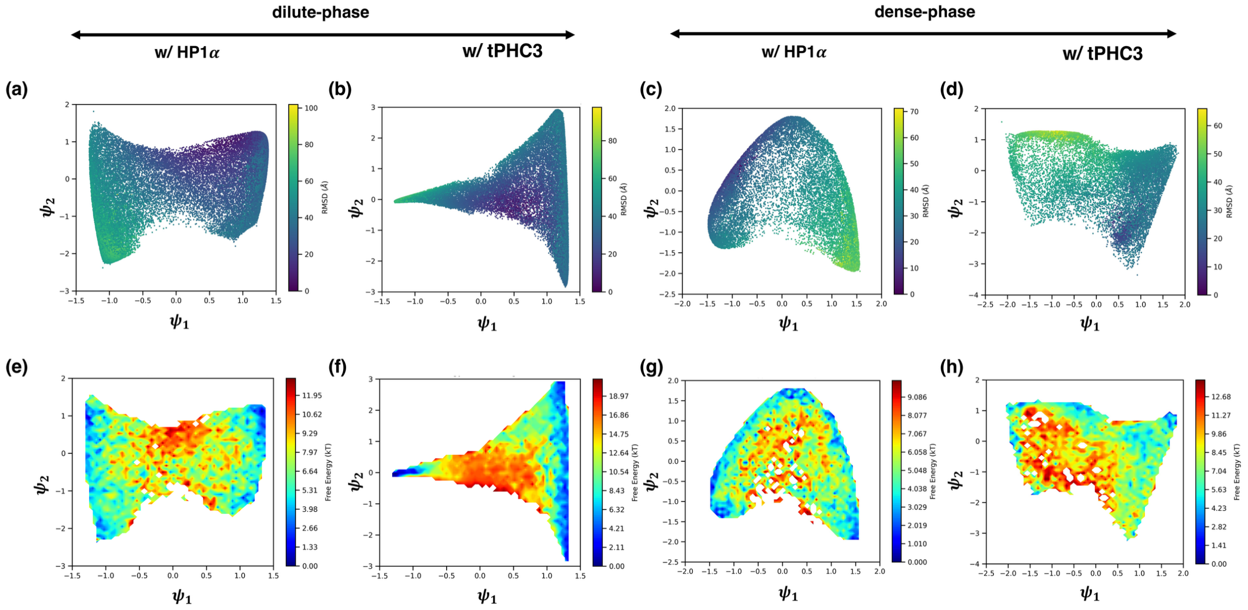


***Figure S5.*** *Sliding window diffusion map analysis of tri-nucleosome substructures across dilute and dense phases with HP1α and tPHC3. (a–d) RMSD-colored projections of the* $\psi_1$*–*$\psi_2$ *diffusion map space constructed from triads of nucleosomes using a sliding window approach. Color scale indicates RMSD (nm) of the tri-nucleosome structure relative to the native α-tetrahedron reference. (e–h) Corresponding free energy surfaces projected onto the same* $\psi_1$*–*$\psi_2$ *coordinates. Each column represents a different simulation condition: (a,e) HP1α in dilute phase; (b,f) tPHC3 in dilute phase; (c,g) HP1α in dense phase; (d,h) tPHC3 in dense phase. In the dilute phase, HP1α (a,e) yields continuous distributions with a modest population of α-tetrahedron–like geometries, while tPHC3 (b,f) displays fragmented and frustrated manifolds with fewer structured intermediates. In the dense phase, HP1α (c,g) sharpens the landscape, enriching compact tri-nucleosome motifs; tPHC3 (d,h) produces a more heterogeneous but enriched ensemble of α-tetrahedron–like intermediates, reflected in multiple low-RMSD basins. These results support a model where tri-nucleosome interactions serve as local intermediates in the global folding of the tetra-nucleosome, with protein identity and phase conditions shaping their stability and prevalence.*

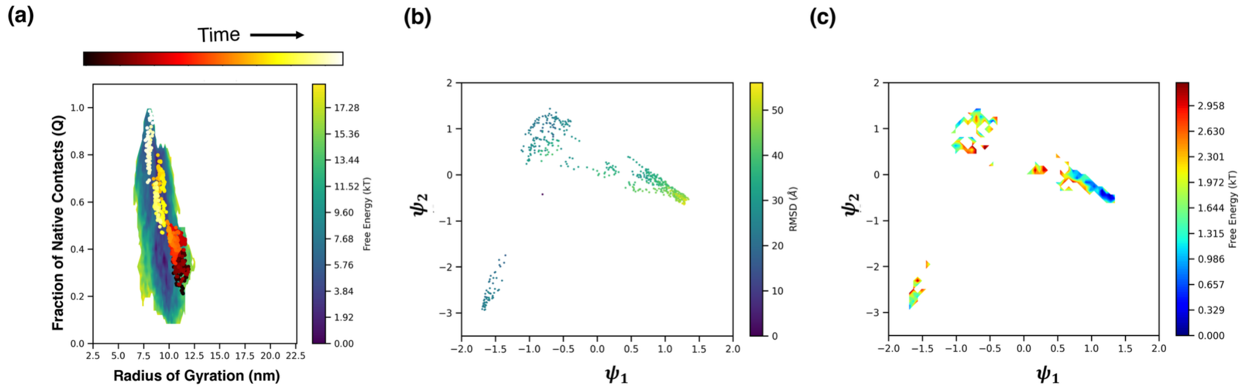


***Figure S6.*** *Reactive folding analysis for the monomeric tPHC3 system. (a) Q–Rg projection of reactive folding subtrajectories colored by simulation time, highlighting folding progression from disordered (low Q, high Rg) to compact (high Q, low Rg) states. (b) Diffusion map embedding ($\psi_1$–$\psi_2$) of the same subtrajectories using RMSD-derived pairwise distances, colored by RMSD from the native crystal structure. Folded states appear as darker green clusters at high $\psi_1$ values, while unfolded conformations are spread across lower RMSD gradients. (c) Corresponding free energy surface projected onto $\psi_1$–$\psi_2$ space. The landscape is fragmented and shallow, with multiple diffuse basins indicating partially folded intermediates and high conformational heterogeneity. Compared to the dimeric system, the monomeric tPHC3 folding path is less structured and more restricted, suggesting that oligomerization enhances topological diversity and folding route accessibility.*

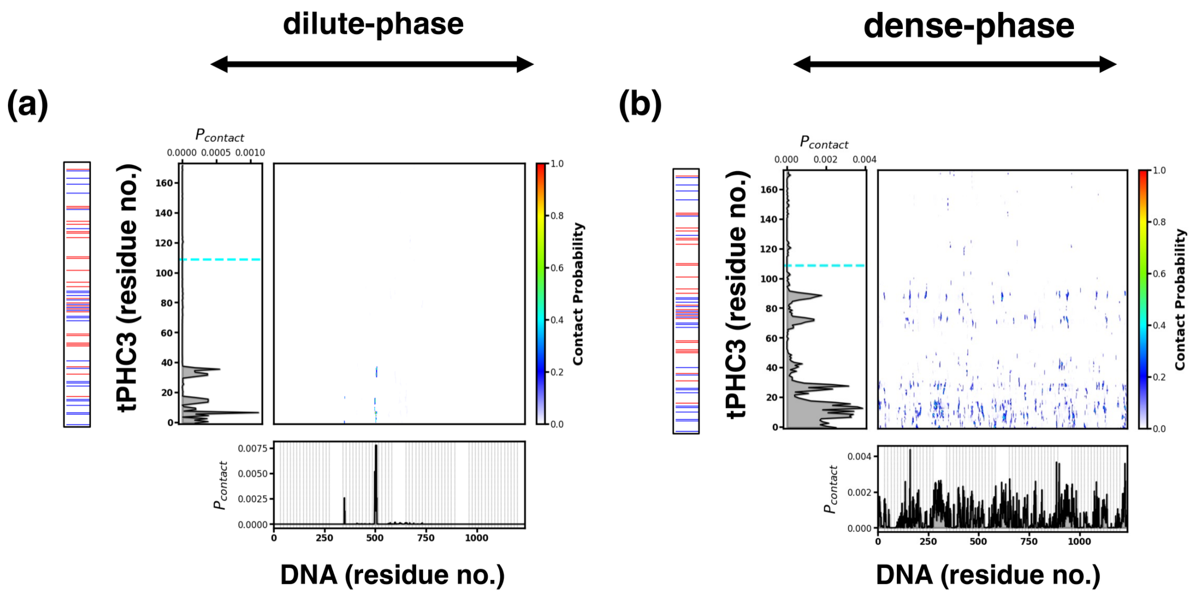


***Figure S7.*** *Protein–DNA contact maps for monomeric tPHC3 in dilute and dense phases. (a, b) Intermolecular contact probability between monomeric tPHC3 and DNA in the dilute (a) and dense (b) phase, computed from WTM-REMD simulations. Vertical lines along the DNA axis indicate superhelical locations on the nucleosomal DNA, with four distinct gaps corresponding to linker regions between the four nucleosomes. Dashed lines along the protein axis delineate the boundaries between different domains of the protein. In the dilute phase (a), tPHC3 exhibits a highly restricted interaction profile, with contacts confined to a narrow segment of the linker DNA and minimal residue diversity. In contrast, the dense-phase system (b) displays widespread interaction across superhelical locations and nucleosomes, though still lacking participation from the SAM domain. These results highlight the enhanced engagement of monomeric tPHC3 with chromatin under condensed conditions and suggest context-dependent remodeling of its binding.*

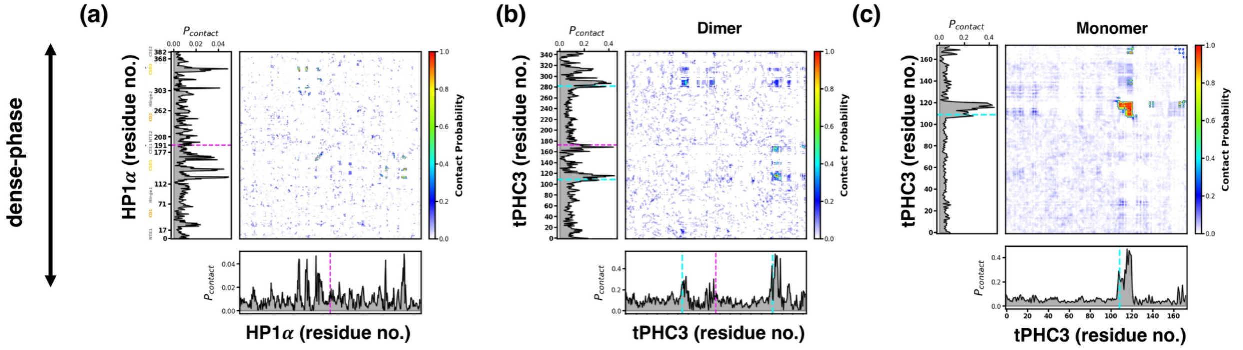


***Figure S8.*** *Intermolecular protein–protein contact maps for HP1α and tPHC3 in the dense phase from WTM-REMD simulations in the presence of tetra-nucleosomes. (a) HP1α shows broadly distributed contacts involving its dimerization interface and disordered regions, consistent with its flexible self-association. (b) Dimeric tPHC3 displays a more focused interaction profile, with strong SAM–SAM and linker-associated contacts, though constrained by pre-formed dimerization. (c) Monomeric tPHC3 reveals emergent SAM–SAM and linker-associated interactions, not observed in the unbiased system, suggesting that chromatin compaction promotes alternative modes of multimerization even in the absence of pre-dimerization. Dotted lines demarcate major domain boundaries.*

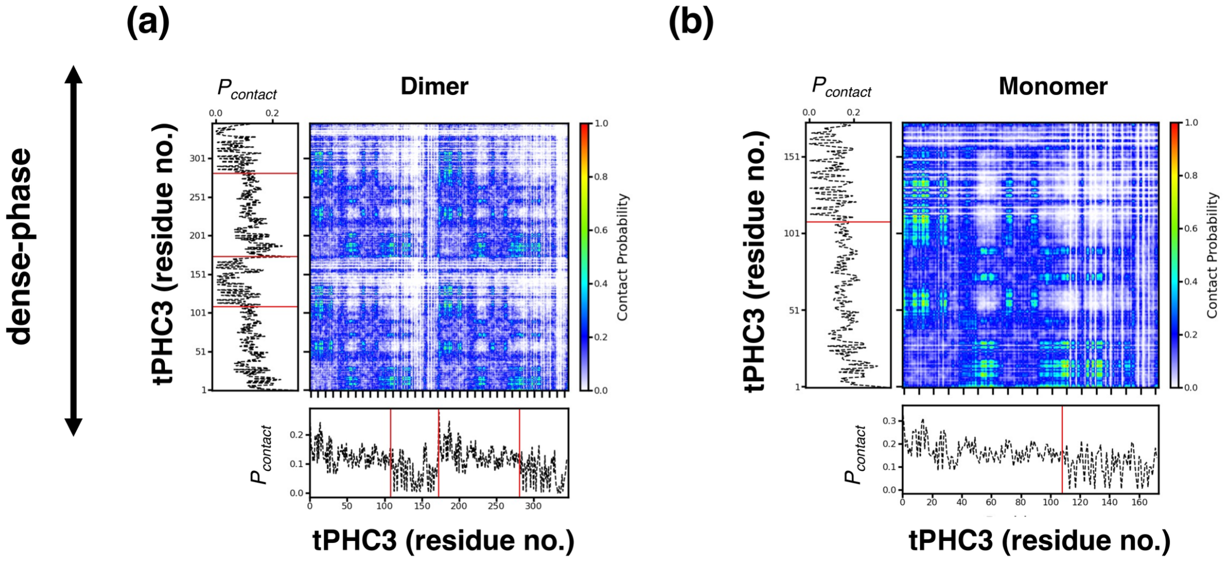


***Figure S9.*** *Intermolecular protein–protein contact maps for dimeric and monomeric tPHC3 from unbiased slab simulations in the absence of nucleosomes (300 K, 100 mM salt). (a) Dimeric tPHC3 exhibits extensive interactions between linker regions and SAM–linker contacts, forming a cohesive self-association network. (b) Monomeric tPHC3 shows similar interaction motifs, including moderate linker–linker and SAM–linker contacts, despite lacking preformed dimerization. These results highlight the intrinsic multimerization propensity of tPHC3, driven by flexible linker-mediated interactions. Red lines indicate domain boundaries.*

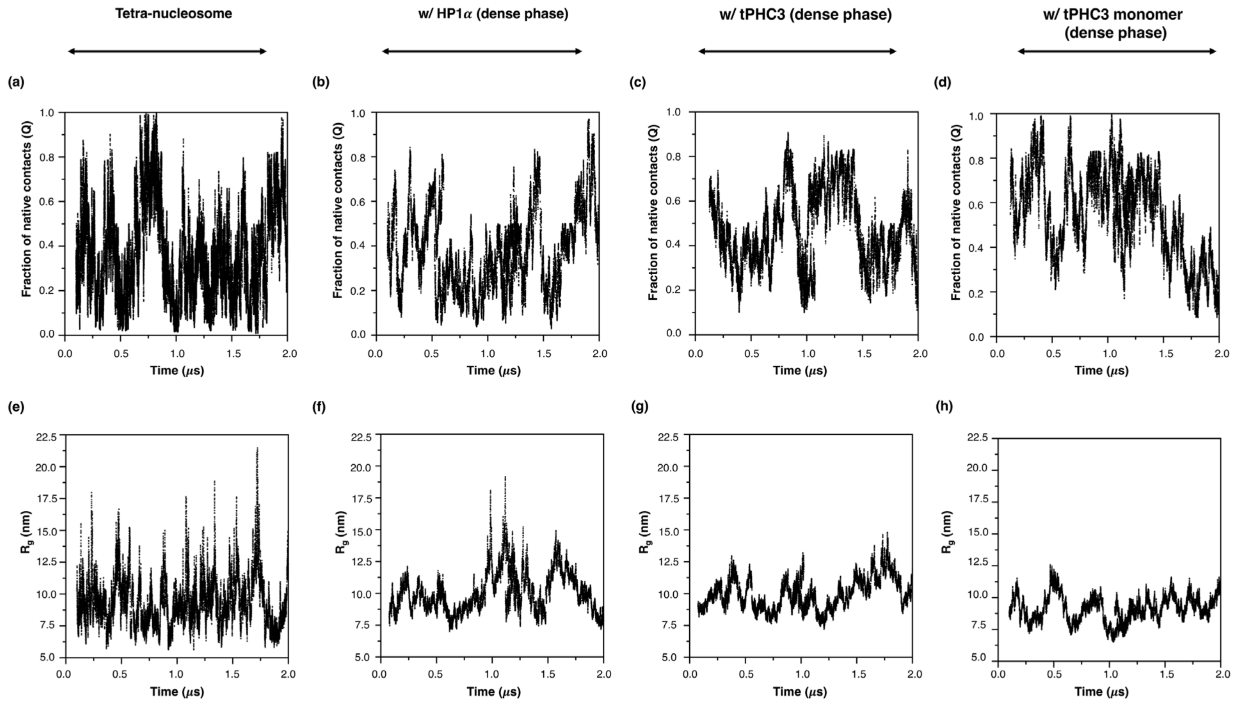


***Figure S10.*** *Sampling of Q (top panel), and $R_g$ (bottom panel) as a function of time at 300 K for protein-free tetra-nucleosome system, and tetra-nucleosome in dense phase conditions of HP1α, tPHC3 in both dimeric and monomeric form. 0.1 μs equilibration data is not included.*

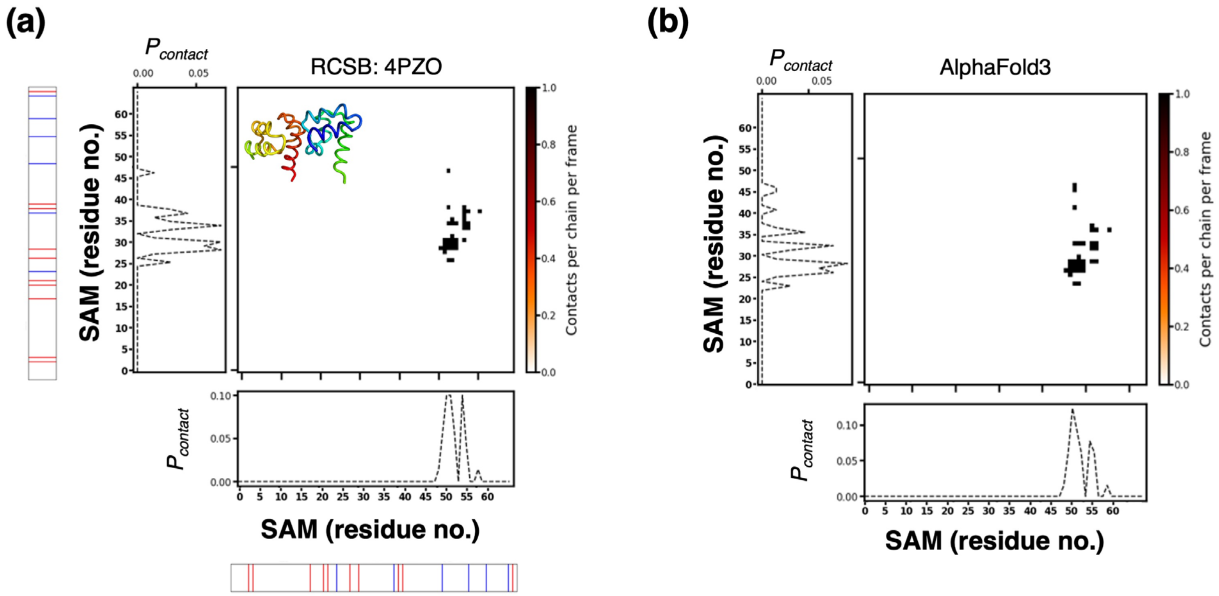


***Figure S11.*** *Reference SAM–SAM contact maps from known structural models. (a) Contact probability map derived from the RCSB crystal structure of the PHC3 SAM domain (PDB ID: 4PZO), showing a well-defined head-to-tail oligomerization interface mediated by charged residues. The inset highlights the corresponding ribbon structure. (b) Contact pattern from an AlphaFold3-predicted SAM dimer, which preserves similar contact geometry, including highly localized interaction hotspots.*
